# Fate plasticity of interneuron specification

**DOI:** 10.1101/2024.10.02.614266

**Authors:** Mohammed A. Mostajo-Radji, Walter R. Mancia Leon, Arnar Breevoort, Jesus Gonzalez-Ferrer, Hunter E. Schweiger, Julian Lehrer, Li Zhou, Matthew T. Schmitz, Yonatan Perez, Tanzila Mukhtar, Julia Chu, Madeline G. Andrews, Frederika N. Sullivan, Dario Tejera, Eric C. Choy, Mercedes F. Paredes, Mircea Teodorescu, Arnold R. Kriegstein, Arturo Alvarez-Buylla, Alex A. Pollen

## Abstract

The generation of neuronal subtypes in the mammalian central nervous system is driven by competing genetic programs. The medial ganglionic eminence (MGE) gives rise to two major cortical interneuron (cIN) populations, marked by Somatostatin (Sst) and Parvalbumin (Pvalb), which develop on different timelines. The extent to which external signals influence these identities remains poorly understood. Pvalb-positive cINs are particularly important for regulating cortical circuits through strong perisomatic inhibition, yet they have been difficult to model *in vitro*. Here we investigated the role of the environment in shaping and maintaining Pvalb cINs. We grafted mouse MGE progenitors into a variety of 2D and 3D co-culture models, including mouse and human cortical, MGE, and thalamic systems with dissociated cells, organoids, organotypic cultures, and conditioned media. Across models, we observed distinct proportions of Sst- and Pvalb-positive cIN descendants. Strikingly, grafting MGE progenitors into 3D human, but not mouse, corticogenesis models led to efficient, non-autonomous differentiation of Pvalb-positive cINs. This differentiation was characterized by upregulation of Pvalb maturation markers, downregulation of Sst-specific markers, and the formation of perineuronal nets. Furthermore, lineage-traced postmitotic Sst-positive cINs, when grafted onto human cortical models, also upregulated Pvalb expression. These results reveal an unexpected level of fate plasticity in MGE-derived cINs, demonstrating that their identities can be dynamically shaped by the surrounding environment.

## Introduction

The cerebral cortex contains both excitatory projection neurons (PNs) and inhibitory interneurons (cINs). While the mechanisms underlying the generation and maintenance of this neuronal diversity remain incompletely understood, recent cell atlas studies reveal a significantly greater diversity of neurons in the adult cortex compared to the prenatal stage. This suggests that external cues during migration and maturation may play a key role in shaping neuronal identity (***Allison et al., 2021***; ***Harb et al., 2022***; ***Cadwell et al., 2019***; ***Di Bella et al., 2021***; ***Nowakowski et al., 2017***; ***Schmitz et al., 2022***; ***Ozair et al., 2018***). During embryonic development, PNs and cINs originate from distinct progenitor pools: PNs are generated locally in the ventricular zone of the developing telencephalon, while cINs migrate long distances from the medial and caudal ganglionic eminences (MGE and CGE) and the preoptic area (POA) to reach their final cortical destinations (***Anderson et al., 1999***; ***Nery et al., 2002***; ***Wichterle et al., 2003***; ***Xu et al., 2004***). Upon reaching the cortex, cINs integrate into local circuits by forming connections with PNs.

Several studies underscore the critical role of extrinsic cues in shaping cIN development (***Wamsley and Fishell, 2017***), influencing migration and lamination (***De Marco García et al., 2011***; ***Lodato et al., 2011***), synaptic wiring (***Wester et al., 2019***; ***Ye et al., 2015***), morphological and electrophysiological maturation (***De Marco García et al., 2011***; ***Pan et al., 2019***), and survival (***Southwell et al., 2012***; ***Wong et al., 2018***; ***Wu et al., 2024***). However, the extent to which external signals specify cIN subtypes remains an open question (***Bragg-Gonzalo et al., 2021***; ***Jabaudon, 2017***; ***Pipicelli et al., 2023***; ***Wu et al., 2024***). For example, findings in a *Fezf2*-knockout mouse model, where subcerebral PNs are replaced by callosal PNs (***Molyneaux et al., 2005***), reveal a fate switch between two sub-classes of Pvalb-positive cINs (***Wu et al., 2024***). Remarkably, even after migration and initial synaptic wiring with PNs, Pvalb/Fzd6-positive cINs lose their molecular subclass identity and instead acquire the identity of Pvalb/Slc39a8-positive cINs (***Wu et al., 2024***). This suggests that extrinsic signals can influence cIN fate well beyond early developmental stages.

Previous studies on MGE-derived cIN progenitors have identified distinct spatial and temporal origins for different cIN subtypes (***Inan et al., 2012***; ***Wonders et al., 2008***; ***Hu et al., 2017a***). For instance, Sst-positive cINs are primarily generated early from dorsal MGE progenitors, whereas Pvalb-positive cINs are mainly produced later from ventral MGE progenitors (***Inan et al., 2012***; ***McKenzie et al., 2019***; ***Wonders et al., 2008***). The transcription factor *Mef2c* has been implicated in early post-mitotic MGE-derived cINs destined to become Pvalb cINs, indicating that early molecular programs guide subtype specification (***Allaway et al., 2021***; ***Mayer et al., 2018***). However, single-cell RNA sequencing of the developing mouse brain shows that cIN progenitors from different ganglionic eminences share similar transcriptional profiles, which diverge later during differentiation (***Allaway et al., 2021***; ***Di Bella et al., 2021***; ***Mayer et al., 2018***; ***Mi et al., 2018***). Furthermore, in contrast to PN subtypes, which follow similar maturation timelines (***Kroon et al., 2019***), MGE-derived cIN subtypes exhibit distinct maturation trajectories: Sst-positive cINs are specified and incorporated into circuits early in development (***Bragg-Gonzalo et al., 2021***; ***Tuncdemir et al., 2016***), whereas Pvalb-positive cINs mature later during postnatal life (***Chattopadhyaya et al., 2004***; ***Hensch, 2005***; ***Ma et al., 2013***). Notably, sensory input plays a critical role in postnatal Pvalb cIN maturation (***Chattopadhyaya et al., 2004***), raising the possibility that early environmental cues may trigger distinct genetic programs that further diversify MGE-derived cIN subtypes.

Grafting studies have been pivotal in uncovering the role of developmental cues in various biological processes (***Alvarado-Mallart, 2000***). For instance, heterochronic transplantation experiments have shown that MGE-derived cINs play an intrinsic role in the maturation of cortical circuits (***Southwell et al., 2010***; ***Tang et al., 2014***). Interspecies chimeric models provide additional insights into how intrinsic and extrinsic factors regulate neuronal maturation rates (***Brown et al., 2021***; ***Linaro et al., 2019***; ***Marchetto et al., 2019***; ***Zheng et al., 2021***). These models are especially valuable because they modify both developmental timing (heterochrony) and gene expression patterns. Human brain development, for example, occurs over a much longer time frame than that of rodents (***Semple et al., 2014***), and many genes have evolved to exhibit different expression patterns across species, even though the underlying cell types are conserved (***Bakken et al., 2021***; ***Hodge et al., 2019***). Differences in timing and gene expression in these interspecies models may help identify novel environmental factors that influence cell fate specification and maturation (***Mostajo-Radji et al., 2020***; ***Wallace and Pollen, 2024***).

Using mouse and human 2D co-culture and 3D organotypic and organoid models, we demonstrate that the cortical environment influences not only the timing of MGE cIN specification but also the final molecular identity of developing cINs. Remarkably, when mouse MGE cells are grafted onto 3D human cortical tissue - both primary organotypic cultures and cortical organoids - they rapidly differentiate a population mainly comprised of Pvalb-positive cINs weeks earlier than in typical development. Additionally, we show that a subset of lineage-traced, early postmitotic Sst-positive cINs can be induced to upregulate Pvalb in the 3D human cortical environment. These findings reveal the plasticity of MGE-derived cIN fate, demonstrate the critical influence of environmental factors in refining neuronal identity, and highlight *in vitro* conditions that promote efficient differentiation of mouse Pvalb cINs.

### Absence of pre-specification in the cIN lineage

To assess the degree of neuronal subtype specification within MGE-derived populations during embryonic development, we analyzed single-cell gene expression data from the developing mouse telencephalon (E13 to P10, see Materials and Methods) (***Bhaduri et al., 2018***; ***Loo et al., 2019***; ***Mayer et al., 2018***; ***Schmitz et al., 2022***). Our analysis identified six postmitotic clusters of cINs originating from *Nkx2.1*-expressing regions in the MGE and ventromedial forebrain (VMF), including three clusters giving rise to striatal cINs and *Npy*-expressing cortical cINs, which indicate early diversification within some MGE-derived populations (***Figure 1***A). However, only a single cluster was identified for the majority of cortical Pvalb and Sst cINs. Cells within this cluster expressed key markers like *Lhx6*, *Maf*, and *Sst*, but did not express *Pvalb* and could not be divided into distinct Pvalb or Sst subtypes at this developmental stage. Instead, subclusters were grouped by differentiation stage, in agreement with previous findings (***Figure 1***B-C) (***Allison et al., 2021***; ***Di Bella et al., 2021***; ***Mayer et al., 2018***; ***Mi et al., 2018***). The genes *ErbB4*, essential for tangential migration to the cortex, and *Mef2c* were expressed by the entire cluster at this stage (***Figure 1***C), though they later exhibit selective reduction in Sst and enrichment in Pvalb cINs (***Allaway et al., 2021***; ***Mayer et al., 2018***; ***Sun et al., 2016***). These findings suggest that during early development, there is limited transcriptomic evidence for the specification of Pvalb subtypes among MGE-derived cortical cINs.

**Figure 1.**
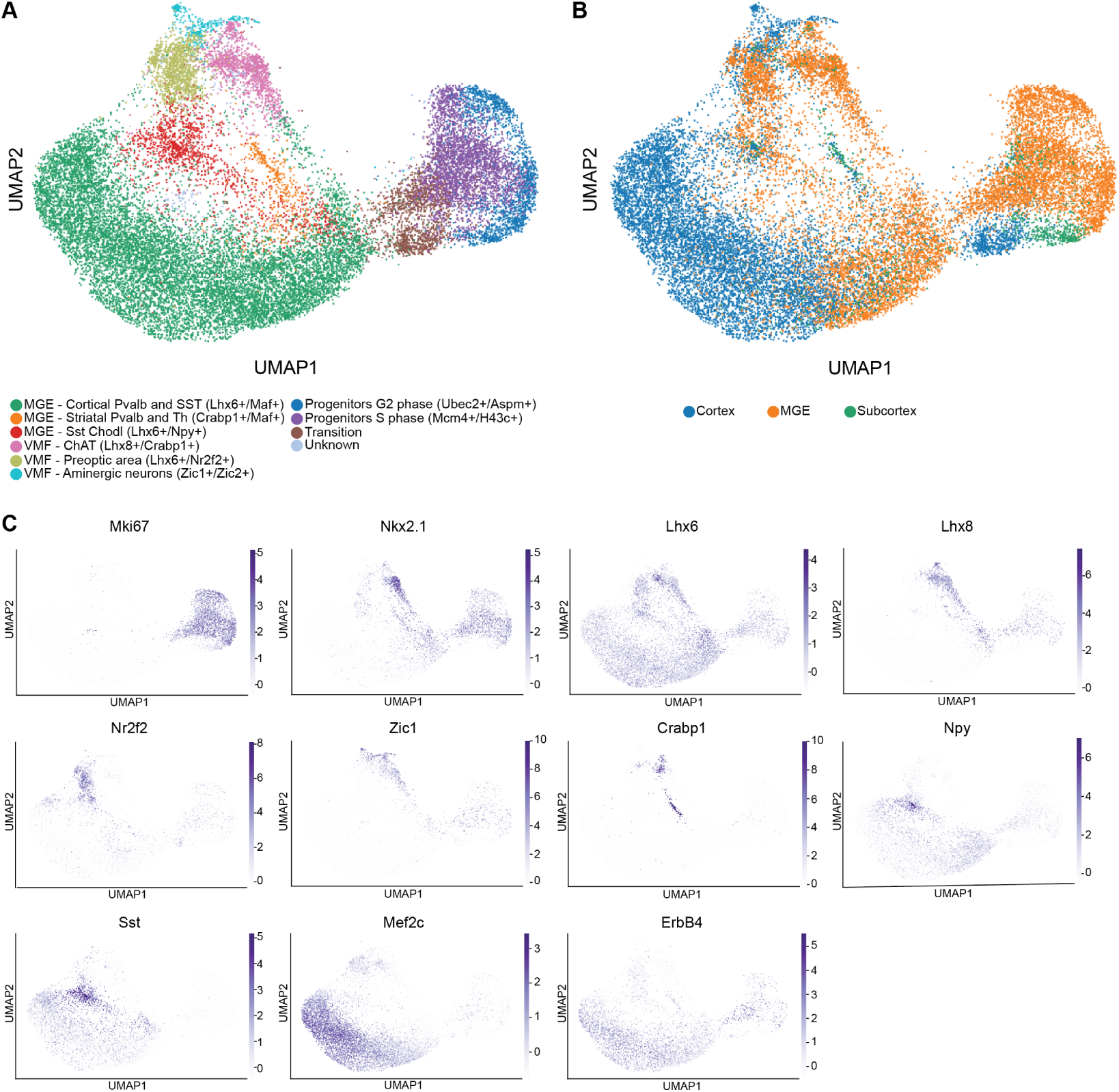
Early diversity of MGE and VMF-derived interneurons. **A.** Single-cell transcriptomic analysis identifies six distinct classes of postmitotic interneurons derived from the MGE and VMF, characterized by the expression of *Gad1*, *Gad2*, *Dlx* genes, *Lhx6*, and *Nkx2.1*. Cells are color-coded by cluster in a UMAP representation of the single-cell transcriptomes. **B.** Classification of cINs based on their region of origin. **C.**

### Human cortical environment induces rapid upregulation of Pvalb in mouse MGE-derived cINs

Previous studies have demonstrated the successful grafting of mouse neuronal progenitors into organotypic brain slices of rats, showing that xenografted mouse neurons can integrate into a host from a different species (***Daviaud et al., 2013***; ***Jäderstad et al., 2010***; ***Tønnesen et al., 2011***). These models have helped clarify cell-cell interactions and the extrinsic regulation of neuronal migration (***Daviaud et al., 2013***). However, because of the relatively similar developmental timelines between these species, the question of whether a heterochronic environment can influence cell fate remains unexplored.

To investigate whether the host environment can influence the identity of neuronal progenitors, we grafted mouse E13.5 MGE-derived cIN progenitors onto organotypic cultures of human and mouse embryonic cortex (***Figure 2***A-C). The donor MGE cells were from the *Nkx2.1*-Cre mouse line, which specifically targets MGE/POA neuronal progenitors and their descendants (***Xu et al., 2008***). We crossed these with the Ai14 mouse line, which contains a floxed td-Tomato reporter gene in the ROSA26 locus (***Madisen et al., 2010***). The identity of the differentiated mouse cINs was determined via immunohistochemistry. Remarkably, the grafted mouse MGE progenitors exhibited distinct fates depending on the host species. Specifically, when grafted onto E14.5 wild-type (WT) mouse brain organotypic slices, 27.00 ± 3.94% of cells differentiated into Sst-positive cINs at 7 days post-graft (DPG), while 72.99 ± 3.94% of the cells were immunonegative for both Sst and Pvalb (***Figure 2***D). Across three grafting batches, none of the cells were Pvalb-positive (***Figure 2***D), consistent with the fact that *Pvalb* is not typically upregulated in mice until the third postnatal week. In contrast, when mouse MGE progenitors were grafted onto human gestational week (GW) 22 primary cortical slices (from three different donors), 82.94 ± 11.60% of the cINs were Pvalb-positive, with 16.81 ± 13.29% remaining double-negative at 7 DPG (***Figure 2***D). Notably, none of the grafted cINs were Sst-positive alone, and only 0.98 ± 1.69% co-expressed both Pvalb and Sst (***Figure 2***D).

**Figure 2.**
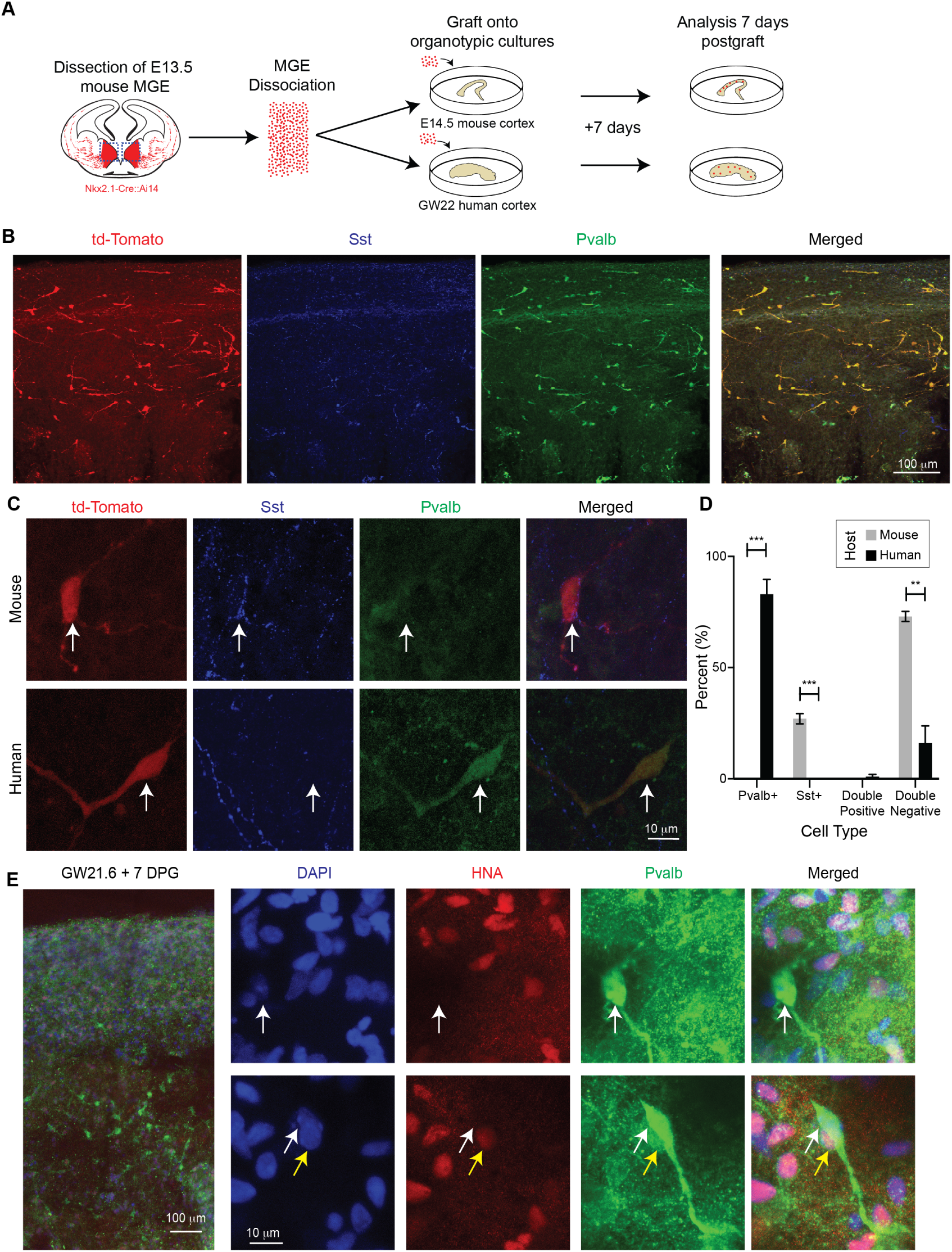
Host-dependent differentiation of mouse MGE progenitors. Grafting of mouse MGE progenitors onto mouse and human cortical organotypic cultures generate different cIN populations at 7 DPG. **A.** Schematic of the experimental design. E13.5 mouse MGE cells were dissociated and grafted onto either GW22 human or E14.5 mouse cortical organotypic cultures to examine cIN differentiation at 7 days DPG. **B.** Representative image of a GW22 human cortical slice grafted with E13.5 td-Tomato-labeled mouse cINs, 7 DPG. **C.** Grafting into E14.5 mouse organotypic cultures predominantly generates Sst-positive cINs, while grafting into GW22 human organotypic cultures primarily induces the differentiation of Pvalb-positive cINs. **D.** Quantification of cIN populations across mouse and human host environments. Statistical significance is indicated as ** = p<0.01 and *** = p<0.001. Error bars represent the standard error of the mean (SEM). **E.** Grafting of unlabeled MGE progenitors allows unbiased identification of Pvalb-positive cINs. White arrows denote mouse nuclei, and yellow arrows point to adjacent human cells. (See additional examples in Supplemental Figure 2). Expression of key marker genes defining each cluster, plotted on the UMAP. Notably, *Pvalb* expression was absent across all analyzed cells.

Importantly, while human MGE-derived cINs migrate into the cortex before GW22 (***Allison et al., 2021***; ***Ma et al., 2013***; ***Zecevic et al., 2011***), resident human cINs do not express Pvalb until postnatal stages (***Allison et al., 2021***; ***Ma et al., 2013***). Consistent with this, we did not detect Pvalb expression in human cells within our organotypic cultures, although Sst-positive cINs in the host tissue were readily observed (***Figure 2***B-C). To verify the functionality of the Pvalb antibody, we performed immunostaining on postmortem samples ranging from GW39 to 57 years old. Pvalb was detected in the GW39 striatum and in older cortical samples, but not in the GW39 cortex, consistent with its developmental absence (Supplemental Figure 1). This confirms that the lack of Pvalb in GW22 organotypic cultures is due to the developmental stage, not an issue with antibody specificity.

To further investigate the observed species-dependent differences in Pvalb induction and minimize potential imaging biases, we conducted an additional experiment in which unlabeled WT cIN progenitors were grafted onto GW21.6-23 cortical slices from three individuals (***Figure 2***D; Supplemental Figure 2). We distinguished mouse and human cells post hoc by immunostaining for human nuclear antigen (HNA), which specifically recognizes human, but not mouse, nuclei. At 7 DPG, we detected Pvalb-positive cINs, all of which were HNA-negative. Notably, Pvalb upregulation occurred at 7 DPG, equivalent to mouse postnatal day 0, demonstrating that the human cortical environment not only induces Pvalb expression in mouse cells but does so at an accelerated rate, three weeks earlier than it would occur *in vivo*.

### Human cortical organoids induce Pvalb expression in mouse cINs: Longitudinal analysis

We next investigated whether transplanting mouse cINs into human cortical organoids could recapitulate the Pvalb induction observed after transplanting into human organotypic slice cultures. Since organotypic cultures are typically limited to about one week (***Croft et al., 2019***; ***Humpel, 2015***), we hypothesized that chimeric induced pluripotent stem cell (iPSC)-derived cortical organoids would allow for long-term analysis of the host environment’s influence (***Kadoshima et al., 2013***; ***Quadrato et al., 2017***). To create a long-term chimeric model, we grafted mouse E13.5 MGE progenitors onto human cortical organoids (***Figure 3***A). Previous studies have demonstrated that deep-layer PNs are crucial for the proper migration of MGE-derived cINs into the cortex during embryonic development (***Lodato et al., 2011***). Thus, we used 6-8 week-old human cortical organoids as hosts, corresponding to the peak period of deep-layer PN neurogenesis (***Pollen et al., 2019***).

**Figure 3.**
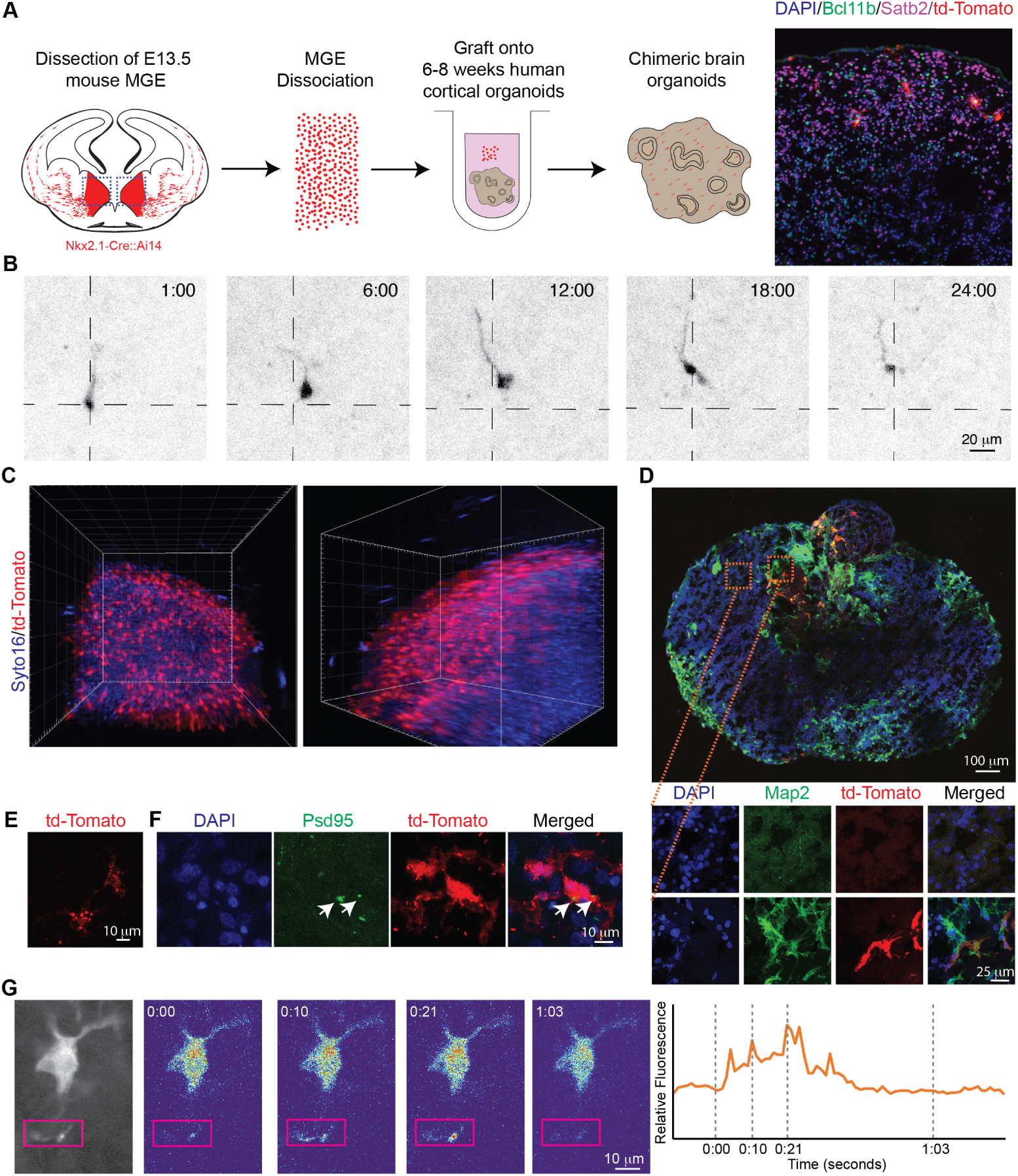
Development of chimeric organoid models. **A.** Experimental design: The MGE from mouse embryos at E13.5 is microdissected and grafted onto human organoids that are 6–8 weeks old. **B.** cIN migration at 1 DPG: Longitudinal live imaging captures a representative migratory cIN over a 24-hour period. The cell soma’s position 1 hour after the experiment starts is marked for reference. **C.** Lightsheet imaging of a grafted organoid. Syto16 dye labels all nuclei within the organoid. **D.** cINs localize to neuronal regions: grafted cINs migrate toward neuronal regions of the organoid, marked by Map2 expression. Top: A whole section of the organoid. Bottom: Comparison of Map2-poor and Map2-rich regions. **E.** Synaptic marker expression at 5 WPG: The *Nkx2.1*-Cre::Ai34 mouse line reveals strong Syp presence in grafted cINs at 5 WPG, indicating the formation of presynaptic vesicles. **F.** Postsynaptic excitatory marker Psd95 puncta are detected in grafted cINs (labeled with *Nkx2.1*-Cre::Ai14) at 5 WPG, suggesting afferent excitatory synapses. **G.** Calcium imaging at 4 MPG: Using the *Nkx2.1*-Cre::Ai96 mouse line, axodendritic calcium transients are observed in grafted cINs (highlighted by magenta rectangles).

To evaluate whether mouse cINs can migrate within the human organoid hosts, we conducted live imaging of chimeric organoids at 1 DPG, capturing hourly observations for 24 hours. Consistent with prior *in vivo* and *in vitro* studies (***Faux et al., 2012***; ***Lepiemme et al., 2021***), we observed that mouse E13.5 MGE cells exhibited high migratory activity within the human cortical organoids. The grafted mouse cells displayed exploratory behavior, neurite branching, and nucleokinesis typical of migrating cINs from the MGE (***Figure 3***B). This indicates that mouse cINs can identify the appropriate substrate and signals within the human cortical environment to support their migration.

To assess the final positioning of the grafted cINs, we employed light-sheet microscopy to visualize whole organoids at 5 WPG (weeks post-graft), after cIN migration and development were complete. We cleared the tissue before imaging to ensure comprehensive visualization of the entire organoid. Our analysis showed that the grafted mouse cINs migrated throughout much of the organoid (***Figure 3***C); however, most were localized at the periphery (***Figure 3***C). In organoids, newborn PNs tend to localize at the periphery, while the center often forms a necrotic core (***Giandomenico et al., 2021***). Several hypotheses could explain the peripheral localization of the grafted cINs: (1) the cINs may be attracted to the periphery, where the majority of excitatory PNs reside, similar to MGE cell migration *in vivo* (***Lodato et al., 2011***); (2) cINs might be unable to penetrate the organoid core; or (3) cINs that migrate to the core may not survive. To differentiate between these possibilities, we used rare organoids in which PNs differentiate internally within the core. After grafting mouse cINs onto these organoids and performing immunostaining for the pan-neuronal marker Map2 at 5 WPG, we consistently found cINs near Map2-positive clusters, irrespective of their location (***Figure 3***D; Supplemental Figure 3A). Notably, Map2-negative regions were largely devoid of mouse cINs, even in peripheral areas (***Figure 3***D; Supplemental Figure 3A). Together, these observations suggest that grafted mouse cINs preferentially localize in areas containing PNs.

Previous studies using monosynaptic tracing of human stem cell-derived PNs grafted into mouse hosts have shown that mouse cINs can form functional synapses with human cortical PNs (***Real et al., 2018***). This suggests that in our grafting paradigm, mouse cINs might integrate into human organoids. To test this, we examined the integration of grafted cINs into human cortical organoids using genetic, molecular, and physiological approaches. First, we used the Ai34 reporter line, which contains a floxed Synaptophysin (*Syp*)-td-Tomato fusion gene inserted into the ROSA26 locus (***Daigle et al., 2018***). Since synaptophysin is a presynaptic vesicle membrane protein, expression of the fusion gene correlates with synapse formation (***Daigle et al., 2018***). At 5 WPG, we detected strong fusion gene expression throughout the grafted cINs, indicating active synapse formation (***Figure 3***E). In complementary experiments, we immunostained grafted *Nkx2.1*-Cre::Ai14 cells for the excitatory postsynaptic marker Psd95 at 5 WPG (***Figure 3***F; Supplemental Figure 3B). Since human PNs are the sole source of excitatory synapses in the organoids, the presence of PSD95 in grafted cINs indicates that the mouse cINs were receiving synaptic input from human PNs.

We next asked whether the grafted cINs exhibited spontaneous activity, a characteristic observed across the central nervous system even in early circuit development (***Allene et al., 2008***). Using calcium imaging, we explored the activity of grafted cINs, leveraging the Ai96 mouse line, which carries the genetically encoded calcium indicator GCaMP6s floxed in the ROSA26 locus (***Madisen et al., 2015***). We crossed these mice with *Nkx2.1*-Cre mice and allowed the cINs to develop for 4 months post-graft (MPG). Calcium imaging in MGE-derived cINs, especially Pvalb-positive cINs, is challenging due to Pvalb’s role as a slow calcium buffer, which dampens calcium transients and complicates imaging (***Caillard et al., 2000***). Therefore, we concentrated our analysis on axodendritic processes, which serve as effective readouts of synaptic integration (***Ali and Kwan, 2019***). At baseline, without external stimuli, we observed strong calcium transients in the axodendritic processes (***Figure 3***G; Supplemental Figure 3C). These results indicate that grafted mouse cINs integrate into human PNs within chimeric organoid models.

### Human cortical organoids recapitulate the accelerated Pvalb expression observed in organotypic cultures

To investigate whether human cortical organoids could recapitulate the accelerated Pvalb expression seen in organotypic cultures, we first analyzed the identity of cINs at 2 DPG, a stage when the cells were still migratory (***Figure 3***B). Strikingly, approximately 50% of grafted cINs were already positive for Pvalb at this early time point (***Figure 4***A; Supplemental Figure 4) (49.51 ± 6.11% Pvalb-positive, 0% Sst-positive, 0.40 ± 0.70% Pvalb and Sst double-positive, and 50.07 ± 6.67% double-negative).

**Figure 4.**
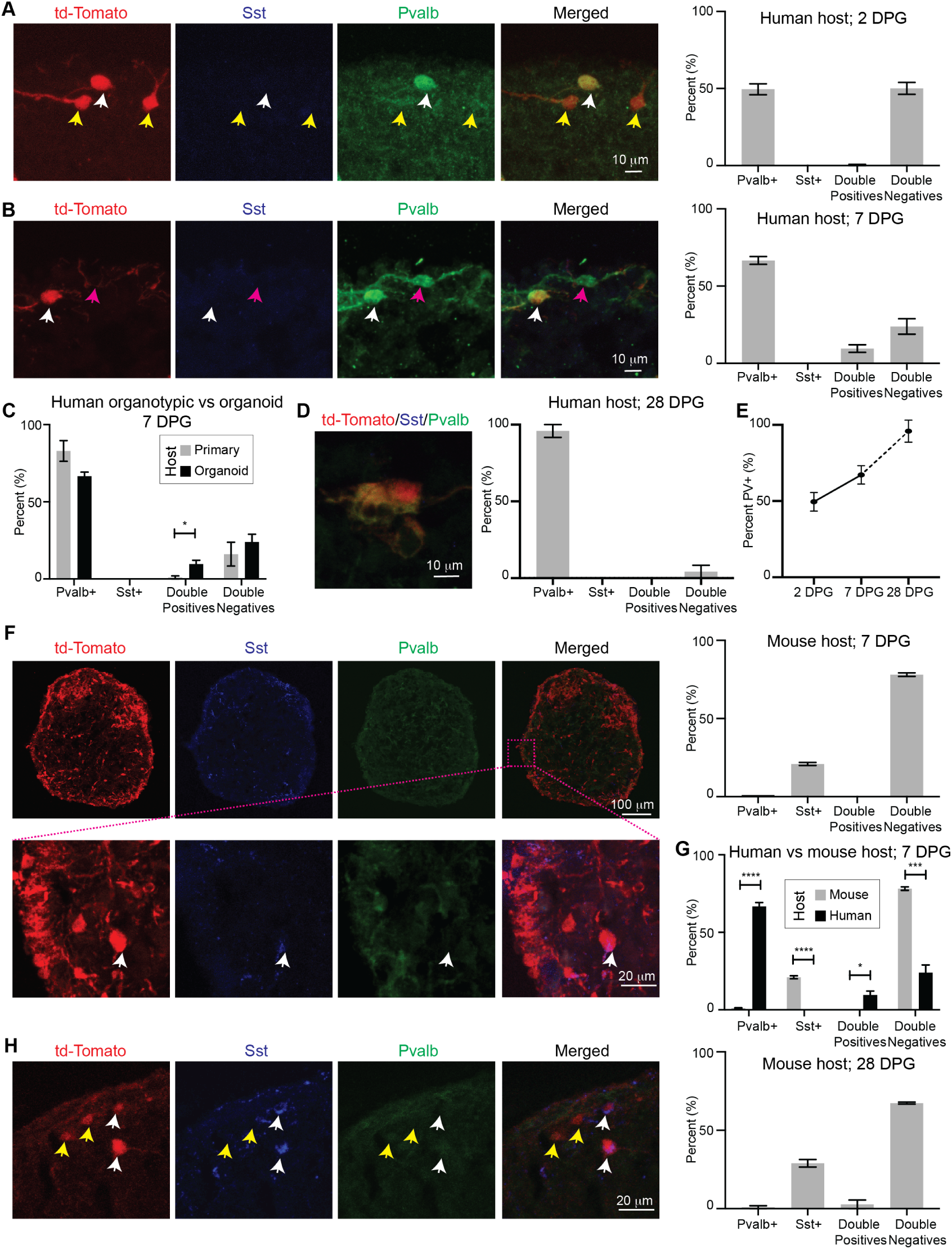
Organoid models recapitulate species-specific biases in Pvalb and Sst expression in grafted cINs. **A-E.** Grafting of mouse cINs onto human organoids: **A.** Representative image and quantification of grafted cINs onto human organoids at 2 DPG. White arrow indicates a Pvalb-positive mouse cIN. Yellow arrows indicate Pvalb-negative mouse cINs. **B.** Representative image and quantification of grafted cINs at 7 DPG. The white arrow indicates a Pvalb-positive mouse cIN, and the magenta arrow indicates a cIN that did not express td-Tomato. **C.** Comparison of cell types generated at 7 DPG between primary human host cultures (gray) and human organoids (black). Data from Figure 2B and Figure 4B are included. **D.** Representative image and quantification of grafted cINs onto human organoids at 28 DPG. **E.** Progressive acquisition of Pvalb expression in cINs over time in human organoid hosts. **F-H.** Grafting of Mouse cINs onto Mouse Organoids: **F.** Representative image and quantification of grafted cINs onto mouse organoids at 7 DPG. **G.** Comparison of cIN populations generated in mouse (gray) versus human (black) organoid hosts at 7 DPG. **H.** Representative image and quantification of grafted cINs onto mouse organoids at 28 DPG. * = p < 0.05; *** = p < 0.001; **** = p < 0.0001. Error bars represent SEM.

We next examined the identity of grafted cINs at later stages. Immunostaining for Pvalb and Sst at 7 DPG revealed that 66.56 ± 4.38% of grafted cINs expressed Pvalb (***Figure 4***B; Supplemental Figure 5), similar to what we observed in mouse cINs grafted onto human GW22 organotypic cultures (***Figure 4***C, p > 0.05). Notably, no cells expressed Sst alone; however, there was a modest but statistically significant increase in double-positive (Pvalb and Sst) cells compared to those grafted onto primary human organotypic cultures (***Figure 4***C, primary human = 0.98 ± 1.69%, organoid = 9.57 ± 4.29%; p < 0.05). The remaining 23.86 ± 8.67% of cINs were negative for both markers, a proportion comparable to that seen in grafts onto organotypic slices (***Figure 4***C, p > 0.05). By 28 DPG, the percentage of Pvalb-positive cINs had increased to 95.83 ± 7.22%, with no Sst-positive or double-positive cells (4.16 ± 7.21% double-negative) (***Figure 4***D). These findings indicate a progressive acquisition of the Pvalb expression in cINs at a faster rate and higher proportion than expected during normal mouse development (***Figure 4***E), suggesting that the 3D human cortical tissue provides extrinsic cues that drive the fate of MGE progenitors.

To independently validate these results, we grafted cINs from *Pvalb*-Cre mice (***Hippenmeyer et al., 2005***) crossed with Ai14 reporter mice. Normally, Pvalb expression in these mice is not detectable until at least three weeks postnatally (***Hensch, 2005***). However, when these *Pvalb*-Cre::Ai14 cINs were grafted onto human cortical organoids, we observed robust td-Tomato expression at 7 DPG, confirmed by immunostaining (Supplemental Figure 6A). This independent approach further supports the accelerated Pvalb expression induced by the human cortical environment.

To assess whether this rapid Pvalb induction is specific to the human cortical environment or can be triggered by any 3D organoid context, we grafted genetically labeled mouse MGE cells onto mouse 3D organoid cultures. We generated mouse primary organoids by dissociating E14.5 mouse cortices and culturing them in neuronal differentiation media (***Ciarpella et al., 2021***), which produces cortical neurons of both upper and deep layer identities (Supplemental Figures 6B-C). In these cultures, only 1.01 ± 0.43% of grafted cINs were Pvalb-positive at 7 DPG. In contrast, 20.91 ± 1.61% of the grafted cINs expressed Sst, while 78.07 ± 1.93% remained negative for both markers (***Figure 4***F-G). These results closely resembled those obtained from grafts onto primary mouse organotypic cultures (***Figure 2***D; Supplemental Figure 6D), suggesting that the mouse cortical environment does not induce accelerated Pvalb expression in mouse MGE cells.

At 28 DPG, a time equivalent to the peak of Pvalb expression *in vivo*, we found that only 0.95 ± 1.65% of grafted cINs in mouse organoids expressed Pvalb (p > 0.05 compared to 7 DPG). However, 28.97 ± 4.18% of the cells were Sst-positive, and 67.30 ± 1.09% remained double-negative (p < 0.01 compared to 7 DPG; Supplemental Figure 6E) (***Figure 4***H). These results, combined with the differences in cIN identities between mouse and human organoids and the progressive acquisition of Pvalb in human hosts (***Figure 4***E; Supplemental Figure 6F), suggest that environmental cues play a critical role in shaping cIN identity.

### 3D Human Environment Promotes Additional Features of Pvalb Identity

Pvalb and Sst are terminal markers of distinct cIN subtypes. Due to the differences in their developmental timelines, additional markers distinguishing these fates have been identified. Notably, the transcription factors *Mef2c* and *Nr2f2* (also known as Coup-TF2) play crucial roles. *Mef2c*, a marker of neuronal maturation (***Pollen et al., 2019***) (Supplemental Figure 7), is thought to identify early Pvalb-fated cINs (***Allaway et al., 2021***; ***Mayer et al., 2018***). In contrast, *Nr2f2* promotes Sst identity and suppresses Pvalb fate within the MGE (***Hu et al., 2017b***). To investigate these markers, we quantified the percentages of Mef2c-positive and Nr2f2-positive cINs at 7 DPG.

Before analyzing the organoids, we validated the Mef2c antibody, which had not been tested in neuronal tissue. We performed immunostaining in the developing cortex at embryonic day 15.5 (E15.5), focusing on the excitatory cortical lineage as an independent and well-characterized system. In this lineage, *Mef2c* labels maturing neurons and is absent in progenitors and newborn neurons (***Di Bella et al., 2021***; ***Nowakowski et al., 2017***; ***Pollen et al., 2019***) (Supplemental Figure 7A). In our validation, we found that Mef2c was excluded from Mki67-positive progenitors in the ventricular and subventricular zones but colocalized with Ctip2-positive, maturing deep-layer PNs in the cortical plate (Supplemental Figure 7B-D). These observations confirmed the specificity of the Mef2c antibody.

We then examined the grafted organoids and found that 90.38 ± 8.99% of mouse MGE-derived cINs grafted onto human organoids upregulated Mef2c at 7 DPG (***Figure 5***A; Supplemental Figure 7E). This result aligns with the observed upregulation of Pvalb in cINs within human 3D cultures. Conversely, no Mef2c-positive cINs were found in grafts onto mouse organoids (p < 0.0001) (*Figure 5*A). In contrast, 25.26 ± 2.19% of cINs grafted onto human organoids were Nr2f2-positive, whereas 90.09 ± 1.50% of cINs in mouse organoids expressed Nr2f2 (p < 0.0001) (***Figure 5***B).

**Figure 5.**
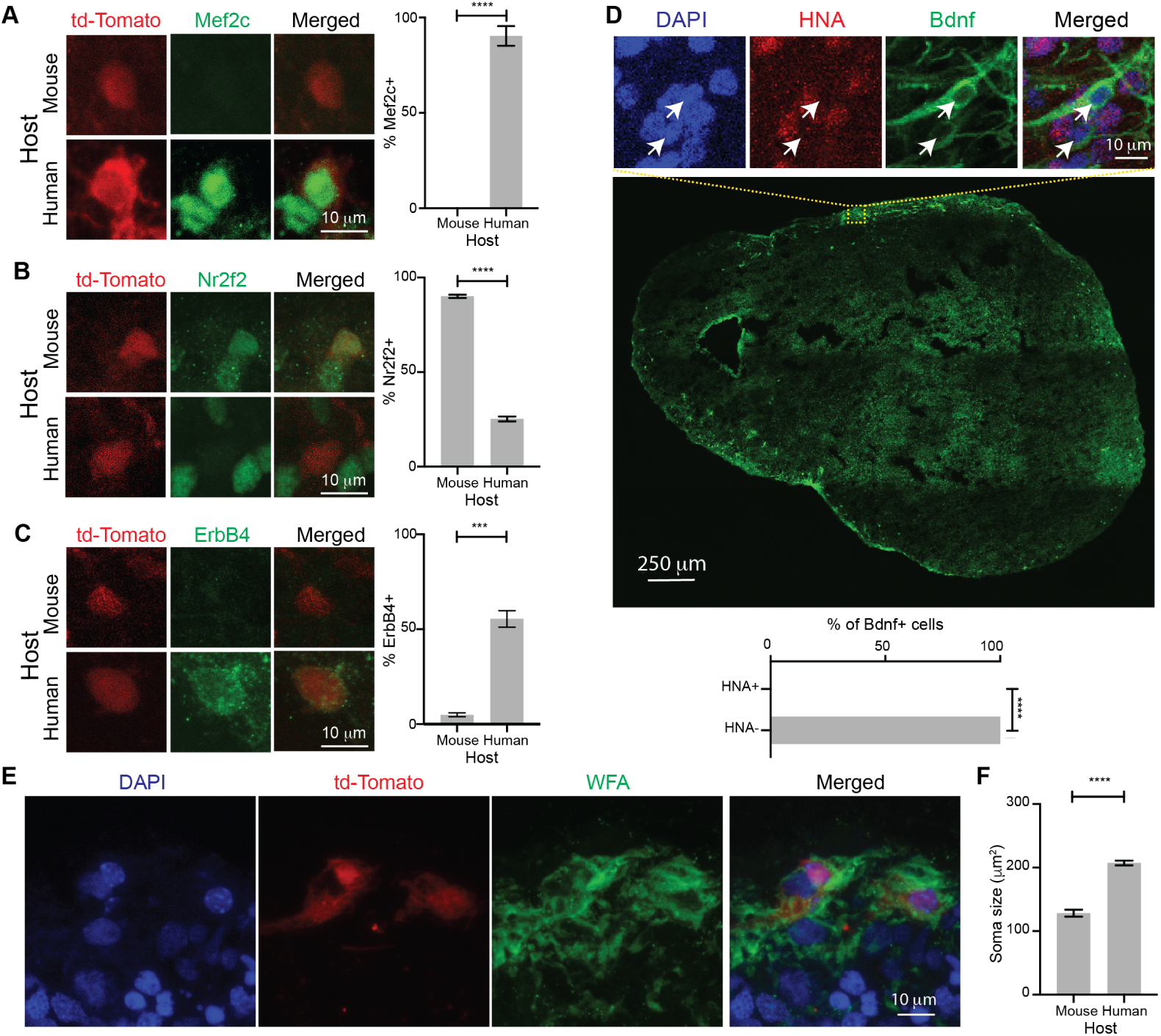
Human cortical organoids influence multiple aspects of Pvalb cIN identity. **A.** Representative images and quantification of Mef2c expression in cINs grafted into mouse and human organoids at 7 DPG. **B.** Representative images and quantification of Nr2f2 expression in cINs grafted into mouse and human organoids at 7 DPG. **C.** Representative images and quantification of ErbB4 expression in cINs grafted into mouse and human organoids at 7 DPG. **D.** Representative image showing Bdnf and HNA staining in human organoids at 8 WPG. All Bdnf-positive cells were HNA-negative, indicating they were not of human origin. **E.** Labeling of PNNs with biotinylated WFA in human organoids at 5 WPG. **F.** Comparison of the soma size of cINs grafted into mouse versus human organoid hosts. *** = p < 0.001; **** = p < 0.0001. Error bars represent SEM.

MGE-derived cells express ErbB4 as they migrate to the cortex, but in mature cortical circuits, ErbB4 is retained exclusively in Pvalb cINs, not Sst cINs (***Sun et al., 2016***). Immunostaining for ErbB4 at 7 DPG again revealed species-specific differences: 55.45 ± 7.49% of cINs grafted onto human organoids expressed ErbB4, while only 4.95 ± 1.76% of cINs grafted onto mouse organoids were positive for this marker (p < 0.001) (***Figure 5***C).

Previous studies showed that a subset of Pvalb-positive cINs also express Bdnf (***Cellerino et al., 1996***; ***Tomas et al., 2020***), while no other cIN subtype is known to produce Bdnf. Bdnf regulates the maturation of Pvalb cINs (***Huang et al., 1999***). Cortical organoids rarely express this gene (***Pollen et al., 2019***; ***Quadrato et al., 2017***; ***Velasco et al., 2019***), and our organoid protocol does not add exogenous Bdnf (Materials and Methods). We therefore performed immunostaining for Bdnf and HNA in human cortical organoids containing mouse cINs at 8 WPG to distinguish the species of the Bdnf-expressing cells. We found extensive Bdnf expression in grafted organoids, but only in HNA-negative (mouse) cells, confirming that the Bdnf-expressing cells were of mouse origin (***Figure 5***D). Next, we analyzed perineuronal nets (PNNs), extracellular matrix structures that preferentially ensheath Pvalb cINs, facilitating their maturation and regulating circuit plasticity (***Wen et al., 2018***). PNNs are influenced by surrounding neurons, including PNs and Pvalb-negative cINs (***Su et al., 2017***). Col19A1, a gene involved in PNN regulation, is highly expressed in mature neurons of cortical organoids (***Pollen et al., 2019***) and in developing deep-layer PNs in the human prefrontal cortex (***Nowakowski et al., 2017***). Using biotin-conjugated Wisteria floribunda agglutinin (WFA), which binds N-acetylgalactosamines in PNNs, we found robust WFA labeling around grafted cINs at 5 WPG. This suggests that the human organoid environment can support PNN formation around grafted Pvalb cINs (***Figure 5***E).

Finally, we measured the soma size of grafted cINs in both mouse and human hosts, as neuronal size is often correlated with identity (***Ye et al., 2015***). Pvalb-positive cINs are among the largest cIN subtypes in the cortex (***Kooijmans et al., 2020***; ***Malik et al., 2019***). We observed that cINs grafted onto mouse organoids had an average soma area of 128.33 ± 65.90 µm², whereas those grafted onto human organoids were significantly larger, with an average soma size of 207.33 ± 39.03 µm² (***Figure 5***F). This is consistent with previous measurements of mouse cIN populations (***Malik et al., 2019***).

In conclusion, alongside the upregulation of Pvalb, Mef2c, ErbB4, and Bdnf, the downregulation of Nr2f2 and Sst, the assembly of PNNs, and the increase in cell size, our data indicate that the 3D human cortical environment induces multiple features of Pvalb identity in mouse-derived cINs.

### Human 3D cortical environment is necessary for Pvalb induction

To determine whether the 3D cortical environment is not only sufficient but also necessary for instructing Pvalb fate in MGE progenitor short-term transplant models, we co-cultured mouse MGE progenitors with dissociated cortical cells from E14.5 mice and GW22 human hosts in a 2D setting. After 7 days in culture (DIC), we analyzed the identity of the grafted cells using immunostaining. Unlike the 3D human environment in organotypic or organoid cultures, no Pvalb-positive cINs were observed in these 2D co-cultures. Furthermore, Sst expression varied depending on the condition: co-culturing with mouse cortical cells resulted in 25.85 ± 4.93% Sst-positive cINs, while co-culturing with human cortical cells led to a 1.7-fold increase, with 40.61 ± 4.94% Sst-positive cINs (p < 0.01) (***Figure 6***A; Supplemental Figure 8A).

**Figure 6.**
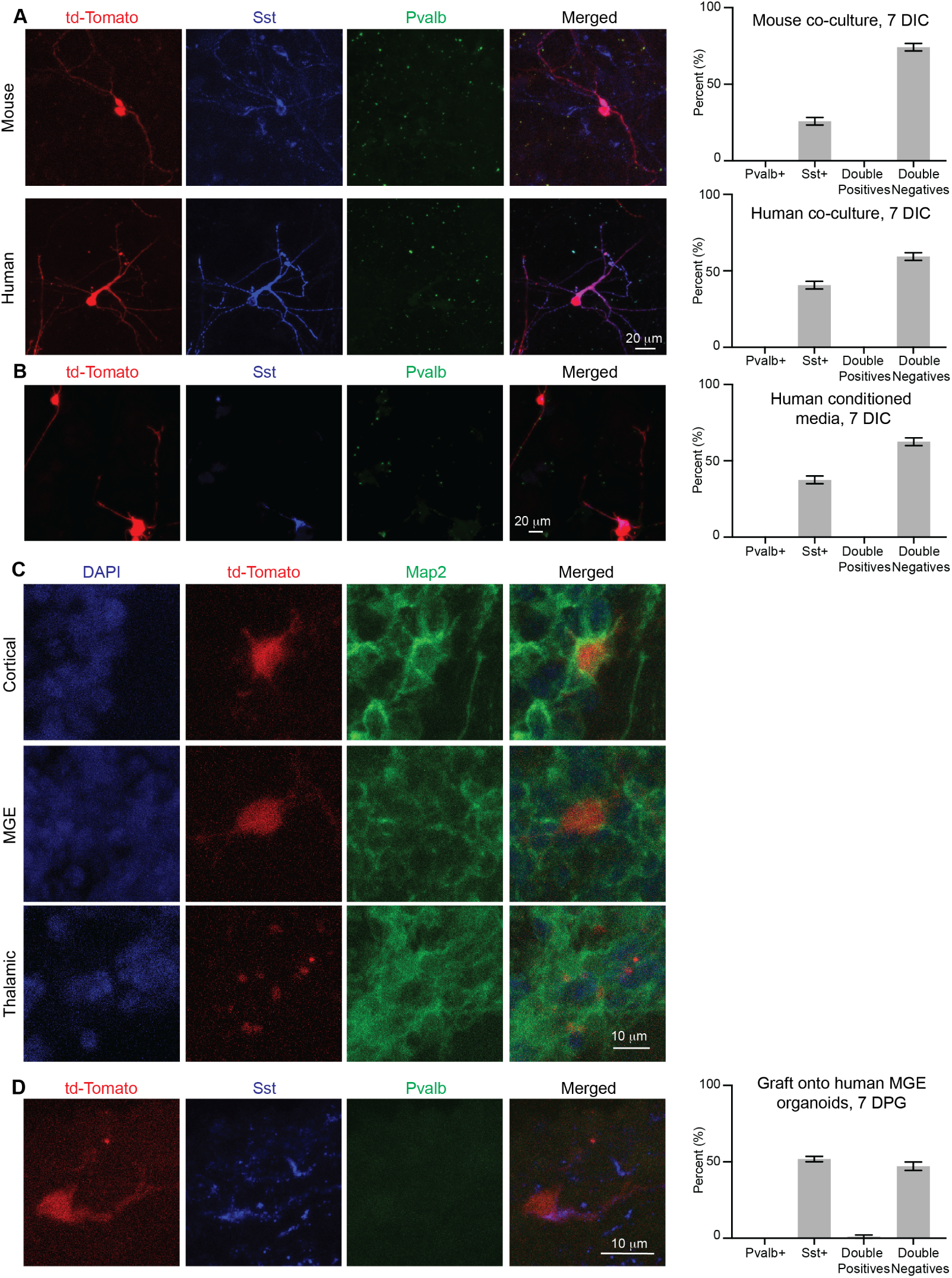
Non-3D human cortical models do not instruct Pvalb fate. **A.** Representative images and quantification of cINs co-cultured with primary mouse and human cortical cells. No Pvalb-positive cINs were observed at 7 DIC. **B.** Representative images and quantification of cINs cultured in media conditioned by primary human cortical cells at 7 DIC. **C.** Representative images of cINs grafted onto cortical, MGE, and thalamic organoids. MAP2, a pan-neuronal marker, is used for reference. **D.** Representative images and quantification of cINs grafted into human MGE organoids at 7 DPG. Notably, additional Sst-positive cells from the host organoid were observed. Error bars represent SEM.

Given that the 2D co-culture with human cortical cells produced a higher proportion of Sst-positive cINs, we questioned whether this increase was driven by direct cell-to-cell interactions or diffusible signals. To explore this, we cultured mouse MGE progenitors in media conditioned by primary human cortical cells grown in 2D. After 7 DIC, the percentage of Sst-positive cINs was similar between cells grown in conditioned media and those co-cultured directly with primary human cells (37.49 ± 5.08% Sst-positive cINs, p > 0.05) (***Figure 6***B; Supplemental Figures 8B-C). This suggests that the Sst induction observed in 2D co-cultures was mediated by diffusible factors in the media.

We then examined whether the specification of Pvalb-positive cINs in our chimeric grafts required a cortical environment, or if Pvalb could be upregulated in other 3D human brain contexts. To address this, we grafted mouse MGE cINs onto human MGE and thalamic organoids. MGE organoids served as a control for matching the regional identity of the cINs’ birthplace, while thalamic organoids represented a nearby region where MGE-derived cINs typically do not migrate (***Jager et al., 2020***) (Supplemental Figure 9A-B). The outcomes were markedly different: at 7 DPG, grafting mouse MGE cINs onto human MGE organoids resulted in 51.78 ± 3.03% Sst-positive cINs, with no cells expressing Pvalb alone. However, 1.07 ± 1.87% of the cells co-expressed both Pvalb and Sst (***Figure 6***C-D; Supplemental Figure 9C-D). In contrast, most cINs grafted onto thalamic organoids died shortly after transplantation (***Figure 6***C). Together, these findings indicate that the 3D human cortical environment is essential for the early induction of Pvalb markers in grafted mouse cINs.

### Human cINs do not upregulate Pvalb shortly after being grafted onto human cortical organoids

Given the rapid upregulation of Pvalb observed in mouse cINs grafted onto human cortical organoids, we next investigated whether grafting human cINs would produce a similar effect. Previous studies involving the fusion of cortical and ventral organoids reported Pvalb upregulation only after several months in co-culture and at low efficiency (***Birey et al., 2017***). This delayed response may be attributed to the slower developmental timeline of human cINs or to missing conditions during iPSC-derived cIN induction that could bias differentiation away from a Pvalb identity.

To minimize potential biases in iPSC induction, we grafted primary cIN progenitors microdissected from the MGE of three different GW18-19 donors onto human cortical organoids (Supplemental Figure 10A). GW18-19 represents the peak of cIN neurogenesis in the human MGE (***Paredes et al., 2022***). The grafted cells were labeled using an adenovirus expressing eGFP under the CMV promoter. Immunolabeling of grafted cells 7 DPG revealed that 99.87 ± 0.34% of eGFP+ cells were negative for both Sst and Pvalb (Supplemental Figures 10B-C). While we cannot rule out Pvalb up-regulation at later time points, we conclude that the early (<7 DPG) Pvalb upregulation is specific to mouse cINs.

### Non-cell-autonomous regulation of Pvalb fate

We next investigated whether non-cell-autonomous factors could influence Pvalb fate acquisition. We first focused on modifying the media composition, specifically the removal of fetal bovine serum (FBS), which is known to contain a complex mixture of nutrients that activate various signaling pathways, including those essential for neuronal maturation (***Bardy et al., 2015***; ***Edwards et al., 2010***; ***Liu et al., 2023***). We hypothesized that removing FBS from the differentiation protocol might affect maturation, including the induction of Pvalb in grafted mouse cINs. To explore this, we generated human cortical organoids under FBS-free conditions and grafted them with mouse cINs (see Materials and Methods). At 45 DIC, we confirmed the presence of corticofugal and early-born callosal PNs in these organoids (Supplemental Figure 11). We analyzed the identity of the cINs at 14 DPG (Supplemental Figures 12-13). Notably, while these cINs expressed several markers associated with Pvalb identity, including *Mef2c, ErbB4,* and the recently identified *Cox6A2*, they maintained an immature phenotype and did not express *Pvalb* itself (Supplemental Figures 12B-F). FBS regulates numerous pathways, including the mammalian target of rapamycin (mTOR) pathway (***Wu et al., 2022***), which has been implicated in the specification of Pvalb-positive cINs (***Amegandjin et al., 2021***; ***Malik et al., 2019***; ***Sharma et al., 2021***; ***Vogt et al., 2015***; ***Wong et al., 2018***; ***Wundrach et al., 2020***). Specifically, activation of the mTOR pathway, such as through the knockout of its upstream inhibitor *Tsc1* in MGE-derived cINs, has been shown to modestly but significantly increase the number of Pvalb-positive cINs in the mouse brain (***Malik et al., 2019***). Therefore, we next assessed the mTOR pathway’s role in Pvalb fate acquisition. We hypothesized that inhibiting mTOR activity with high concentrations (250 nM) of rapamycin would reduce the number of Pvalb-expressing cINs in human cortical organoids. To test this, organoids cultured under the original conditions (with FBS) were treated with rapamycin for 14 days starting on the day of grafting, while control organoids received vehicle treatment (***Figure 7***A). mTOR activity was assessed by immunostaining for phosphorylated ribosomal protein S6 (pS6), a downstream marker of mTOR signaling and a well-established indicator of Pvalb fate in MGE-derived cINs (***Malik et al., 2019***). In the control group, 90.00 ± 8.82% of grafted cINs were pS6-positive at 14 days DPG (***Figure 7***B), consistent with earlier results (***Figure 4*** and ***Figure 5***). Surprisingly, 85.78 ± 12.84% of rapamycin-treated grafted cINs also showed pS6 positivity (p > 0.01) (***Figure 7***B), indicating that phosphorylation of S6 in grafted MGE-derived cINs was resistant to rapamycin inhibition.

**Figure 7.**
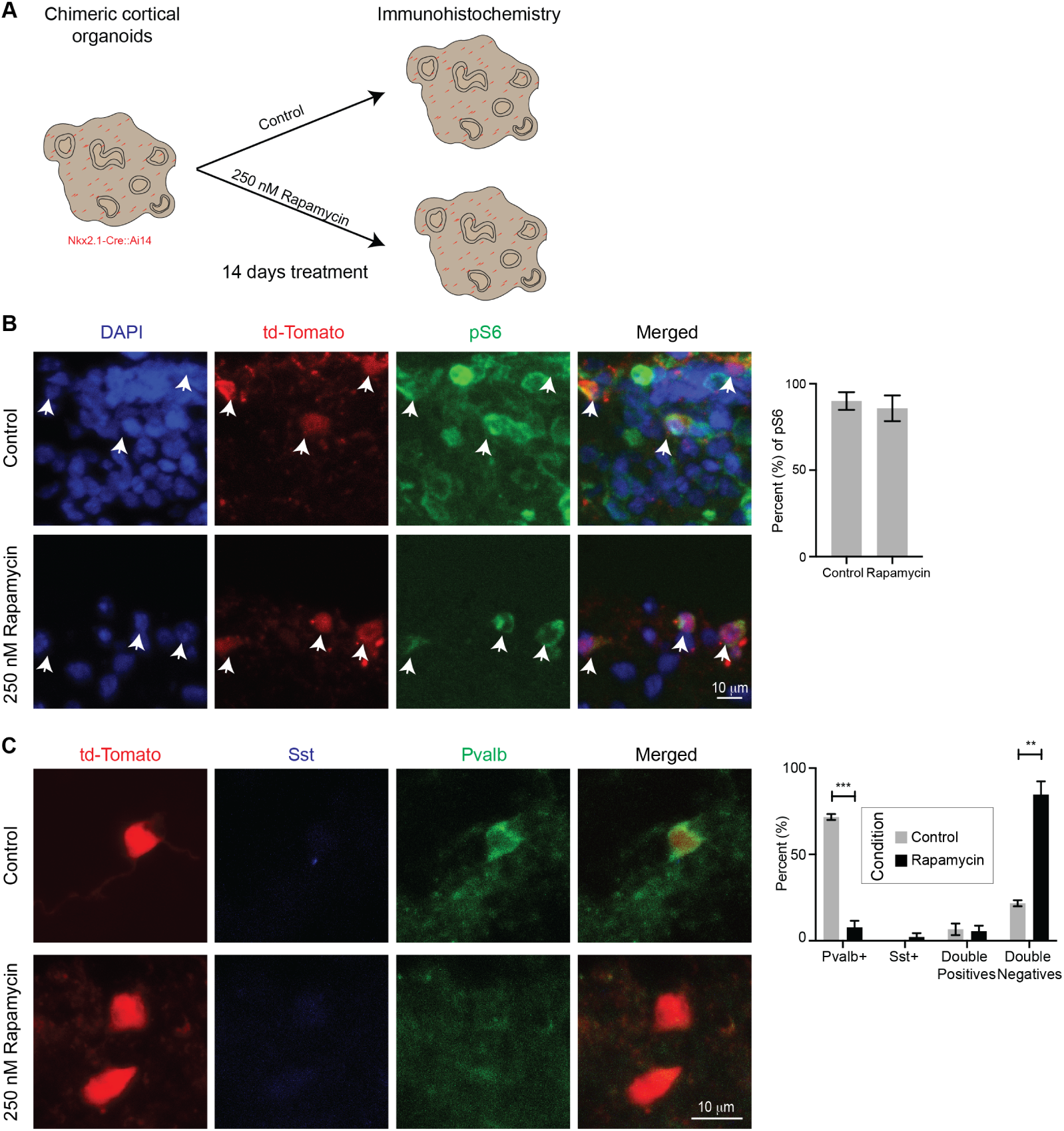
Inhibition of the mTOR pathway in PNs reduces Pvalb specification. **A.** Experimental design: Chimeric cortical organoids were treated with either 250 nM Rapamycin or a vehicle (control) starting at the time of grafting and continuing for 14 days. **B.** Representative images and quantification of phosphorylated ribosomal protein S6 (pS6) in both control and Rapamycin-treated organoids at 14 DPG. White arrows indicate cINs positive for both td-Tomato and pS6. **C.** Comparison of cIN subtypes in control versus Rapamycin-treated organoids. ** = p < 0.01; *** = p < 0.001. Error bars represent SEM.

Typically, ERK/Mapk activity is low in cINs compared to PNs (***Holter et al., 2021***). The rapamycin-resistant pS6 expression observed in the grafted cINs could potentially result from upregulation of the ERK/Mapk pathway (***Roux et al., 2007***; ***Sawicka et al., 2016***). Prior studies have shown that hyperactivation of ERK/Mapk in the MGE selectively reduces Pvalb-positive cINs while leaving Sst-positive cINs unaffected (***Holter et al., 2021***). To explore this possibility, we quantified phosphorylated ERK1/2 (pERK) levels in both control and rapamycin-treated organoids at 14 DPG. The overall percentage of cINs expressing pERK was low and did not differ significantly between control and rapamycin-treated groups (11.69 ± 4.36% in controls vs. 14.04 ± 4.35% in rapamycin-treated organoids, p > 0.05) (Supplemental Figure 14A). These findings suggest that the observed pS6 expression is not due to aberrant ERK/Mapk activation following rapamycin treatment.

The developing human cortex is known to have high mTOR activity in the PN lineage (***Andrews et al., 2020***; ***Nowakowski et al., 2017***), a pattern also observed in organoid models (***Andrews et al., 2020***; ***Pollen et al., 2019***). Consistent with our previous work (***Andrews et al., 2020***; ***Pollen et al., 2019***), we found that rapamycin treatment nearly eliminated pS6 expression in the host cells, with only 1.82 ± 1.34% of rapamycin-treated cells showing pS6 positivity, compared to 31.39 ± 9.39% in controls (p < 0.01) (Supplemental Figure 14B). It is well established that mTOR inhibition in the PN lineage affects the morphology and connectivity of postmitotic PNs (***Chow et al., 2009***; ***LiCausi and Hartman, 2018***). This differential response - grafted cINs being resistant to rapamycin while host PNs were sensitive - allowed us to examine the impact of PN lineage manipulation on the acquisition of Pvalb fate in grafted cINs. At 14 DPG, 71.66 ± 2.91% of grafted cINs in control organoids were Pvalb-positive, consistent with our earlier observation of progressive Pvalb fate acquisition (***Figure 4***E). However, in rapamycin-treated organoids, only 7.69 ± 6.67% of grafted cINs expressed Pvalb (p < 0.001), with 84.62 ± 13.34% of cINs negative for both Pvalb and Sst (***Figure 7***C). These results suggest that mTOR inhibition in PNs significantly influences Pvalb specification in cINs. In conclusion, our findings suggest that Pvalb fate specification is governed by non-cell-autonomous mechanisms that depend on mTOR activity in host cells.

### Early postmitotic Sst-positive cINs can be induced to upregulate Pvalb

*In vivo* lineage reprogramming experiments in mouse cortical PNs have shown a progressive loss of fate plasticity as neurons develop and mature, with a sharp decline shortly after neurons become postmitotic (***De la Rossa et al., 2013***; ***Rouaux and Arlotta, 2013***; ***Ye et al., 2015***). However, whether similar mechanisms to preserve cell fate occur in cINs and whether these mechanisms are intrinsic and universal throughout the central nervous system remains unclear (***Amamoto and Arlotta, 2014***). To address this, we asked whether postmitotic, lineage-traced Sst-positive cINs could be induced to upregulate Pvalb after grafting into a human cortical organoid.

We utilized the well-characterized *Sst*-Cre mouse, which expresses Cre recombinase specifically in postmitotic Sst cINs with minimal leakage into other cIN subtypes (***Malik et al., 2019***; ***Taniguchi et al., 2011***). By crossing this mouse with the Ai14 reporter mouse, we were able to label postmitotic Sst cINs during embryonic development (***Malik et al., 2019***; ***Taniguchi et al., 2011***). We modified our protocol to enrich this population (***Figure 8***A): (1) Dissections were performed at E14.5, when Sst-positive cINs have acquired their identity (***Inan et al., 2012***); and (2) as postmitotic cINs are highly migratory, we dissected the entire ventral telencephalon to account for migrating cINs en route to the cortex.

**Figure 8.**
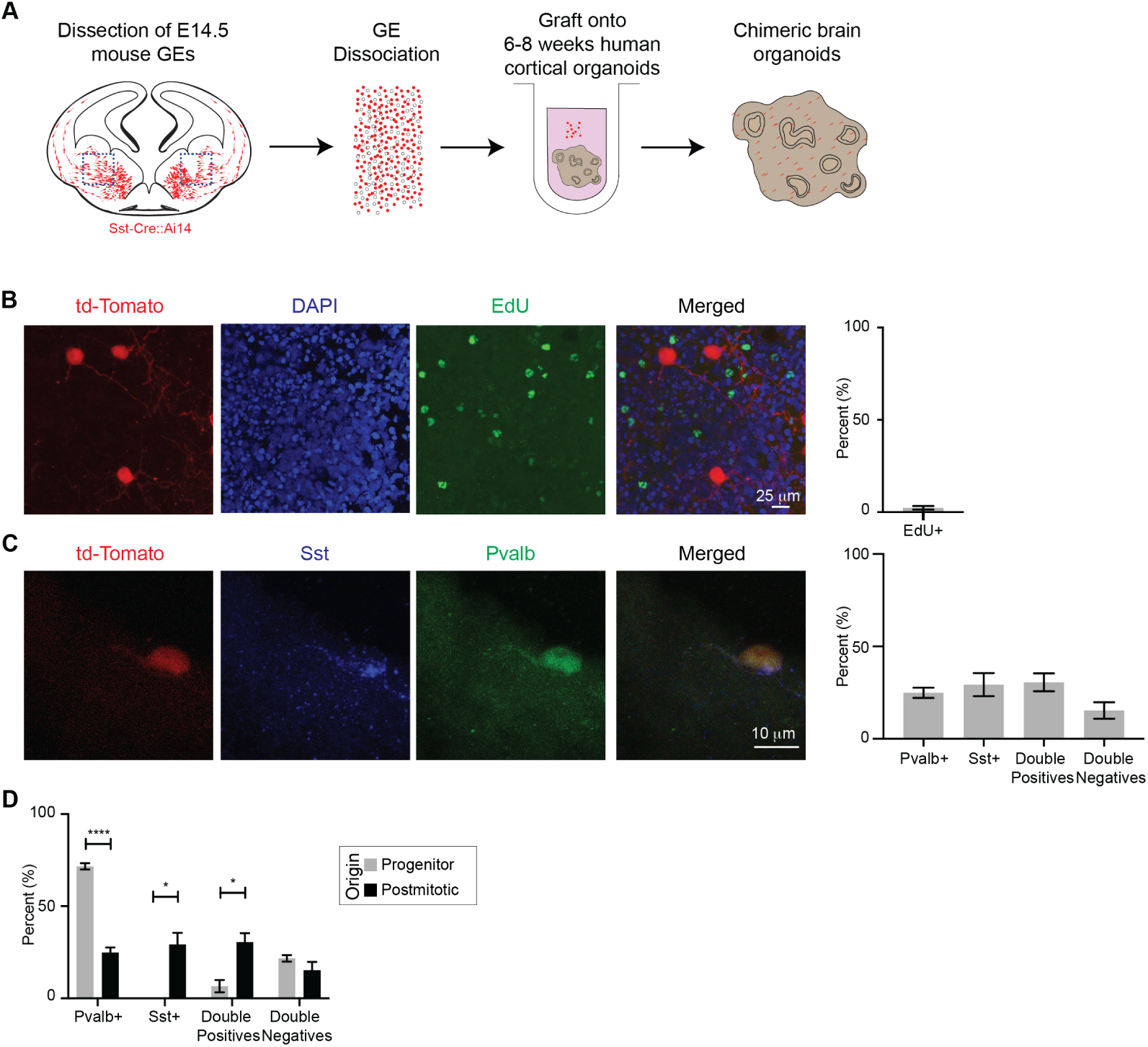
Induction of Pvalb in early postmitotic Sst-positive cINs. **A.** Experimental design: The entire ventral telencephalon was dissected, dissociated into single cells, and grafted onto human brain organoids. **B.** Representative image and quantification from an EdU incorporation assay, showing the postmitotic state of grafted cINs. **C.** Representative image and quantification of the identity of grafted cINs at 14 DPG in human organoids. **D.** Comparison of cell types generated at 14 DPG in organoids grafted with either E13.5 *Nkx2.1*-Cre::Ai14 (Progenitor origin) or E14.5 *Sst*-Cre::Ai14 (Postmitotic origin) grafts. * = p < 0.05; **** = p < 0.0001. Error bars represent SEM.

To confirm the postmitotic state of the grafted cells, we treated them with 5-ethynyl-2’-deoxyuridine (EdU) immediately after grafting and for 2 days. EdU, a thymidine analog, is incorporated into the DNA of dividing cells. Analysis of EdU incorporation at 14 DPG confirmed that only 2.31 ± 3.05% of grafted cells were EdU-positive, indicating that the vast majority of *Sst*-Cre::Ai14 cells were postmitotic at the time of grafting (***Figure 8***B).

Next, we performed immunostaining for Pvalb and Sst at 14 DPG (***Figure 8***C-D; Supplemental Figure 15). We observed that only about one-third (29.28 ± 13.99%) of the grafted *Sst*-Cre::Ai14 cINs were positive for Sst alone (***Figure 8***C). Conversely, half of the grafted cINs were immunopositive for Pvalb, with 24.89 ± 6.18% positive for Pvalb alone, and 30.51 ± 10.79% double-positive for both Pvalb and Sst (***Figure 8***C).

Considering the distinct populations of Sst-positive and Pvalb-positive cINs derived from either MGE progenitors or postmitotic Sst-positive cINs (***Figure 8***D; ***Figure 9***), we conclude that cINs undergo a progressive restriction of cell fate plasticity. Remarkably, however, over half of the grafted postmitotic Sst-positive cINs were still capable of upregulating Pvalb, despite having already expressed Sst. Given that leakage of the *Sst*-Cre mouse into Pvalb cINs is less than 10% (***Malik et al., 2019***) and that the co-localization of Pvalb and Sst is rare in the cerebral cortex (***Gonchar et al., 2008***; ***Malik et al., 2019***; ***Rudy et al., 2010***), these results reveal an unexpected level of plasticity in the fate of cortical cINs.

**Figure 9.**
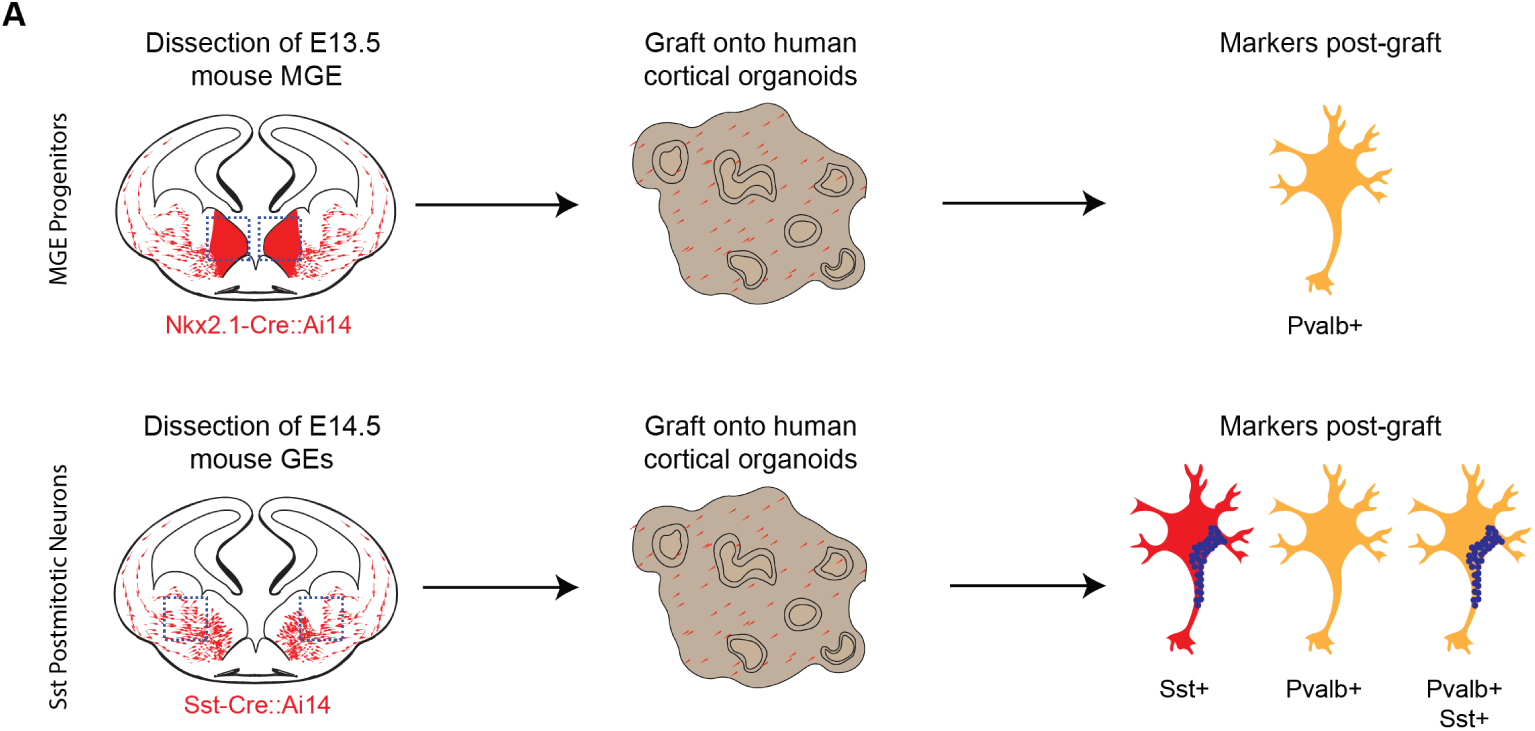
Comparison of Pvalb and Sst upregulation in progenitor vs. postmitotic cIN grafts. **A.** Schematic representation of the experiments. Top: Grafting mouse E13.5 cIN progenitors into human cortical organoids results in the upregulation of Pvalb in the grafted cells. Bottom: Grafting mouse E14.5 lineage-traced Sst cINs results in cINs expressing either Sst, Pvalb, or co-expressing both markers (Sst/Pvalb double-positive). Colors and patterns are reflective of the immunostainings. Note: In both experiments, a minority of cINs were double-negative for Sst and Pvalb.

## Discussion

*In vitro* studies have offered valuable insights into the genetic mechanisms regulating neuronal differentiation. For example, both primary and ES-derived cortical progenitors cultured *in vitro* have been shown to generate PNs in a sequential manner, with deep-layer neurons emerging before upper-layer neurons, closely mimicking *in vivo* development (***Gaspard et al., 2009***; ***Shen et al., 2006***). Moreover, ES-derived PNs retain their areal identity, and when grafted, they selectively integrate into cortical regions that match their origin (***Espuny-Camacho et al., 2018***). These observations suggest that intrinsic factors predominantly control the specification of PN subtypes. Unlike PNs, MGE progenitors and Sst-positive cINs have been successfully differentiated from pluripotent stem cells in both 2D and 3D cultures. However, Pvalb-positive cINs are observed at much lower frequencies *in vitro* (***Allison et al., 2021***; ***Bagley et al., 2017***; ***Birey et al., 2017***; ***Liu et al., 2013***; ***Maroof et al., 2013***; ***Nicholas et al., 2013***; ***Samarasinghe et al., 2021***; ***Sharf et al., 2022***; ***Xiang et al., 2017***). The percentage of Pvalb-positive cINs can be increased by co-culturing with PNs over extended periods (***Allison et al., 2021***; ***Bagley et al., 2017***; ***Birey et al., 2017***; ***Liu et al., 2013***; ***Maroof et al., 2013***), suggesting that interactions with PNs may contribute to Pvalb identity specification. Consistent with this, we observed that grafting mouse MGE progenitors onto human MGE organoids did not induce Pvalb expression (***Figure 6***), whereas grafting them into human cortical hosts led to a significant upregulation of Pvalb (***Figure 2*** and ***Figure 4***). These results suggest that the local environment is a critical determinant of cIN identity.

Modulation of Pvalb identity has been previously observed in heterochronic transplantation studies. However, the appearance of Pvalb-positive cells was delayed, and the proportion of Pvalb-positive cINs was lower than expected. For example, when MGE progenitors were grafted into the visual cortex of juvenile mice, the majority differentiated into Sst-positive cINs, with only 30% adopting a Pvalb fate after one month (***Southwell et al., 2010***). Grafting the same progenitors into the adult mouse visual cortex slightly increased the proportion of Pvalb-positive cINs to 40%, while 20% became Sst-positive (***Davis et al., 2015***). Notably, grafting MGE progenitors into the adult hippocampus reduced Pvalb expression to 10%, with most cells adopting an Sst fate (***Hunt et al., 2013***). These results suggest a previously underappreciated degree of fate plasticity in MGE progenitors, which is influenced by the surrounding environment.

In this study, we identified environmental conditions that promote rapid and extensive differentiation of MGE progenitors into Pvalb-positive cINs. These cINs developed further characteristics of Pvalb identity, including the expression of additional molecular markers, the loss of Sst markers, an increase in soma size, and the formation of PNNs surrounding their soma. Furthermore, we showed that approximately 50% of lineage-traced, early postmitotic Sst-positive cINs can upregulate Pvalb when grafted into a human cortical environment (***Figure 8***). This finding is reminiscent of the PN lineage, where overexpression of *Fezf2* in early postmitotic callosal PNs induced corticofugal molecular programs in 50% of the overexpressing cells (***Rouaux and Arlotta, 2013***; ***Ye et al., 2015***). However, in contrast to PNs, where *Fezf2* fails to alter fate in adult neurons (***Rouaux and Arlotta, 2013***), MGE-derived postmitotic cINs may retain plasticity in their differentiation potential into adulthood (***Dehorter et al., 2015***). Fate plasticity has also been suggested in other interneuron populations. For example, sensory input has been shown to induce a neurotransmitter switch between Sst-positive and dopaminergic interneurons in the adult hypothalamus (***Dulcis et al., 2013***; ***Meng et al., 2018***), and a similar transition has been speculated to occur in primate cINs (***Doll et al., 2024***; ***Ma et al., 2022***; ***Sousa et al., 2017***). Additionally, increased colocalization of the CGE-associated protein calbindin and Pvalb in humans has been proposed as a cortical specialization (***Del Rio and De Felipe, 1997***). Furthermore, scRNA-seq data have revealed extensive colocalization of neurotransmitters in cINs across species and transcriptional similarities between cortical Pvalb and Sst cINs (***Avalos and Sprecher, 2021***; ***Lee et al., 2023***; ***Yao et al., 2021***). However, the fast and direct transition between these cIN cell types has not been previously shown.

It is generally believed that neurons of the central nervous system possess an intrinsic maturation clock that resists acceleration by external factors, such as developmental cues or the electrophysiological properties of cortical PNs (***Barry et al., 2017***; ***Linaro et al., 2019***; ***Marchetto et al., 2019***). However, it has been shown that Pvalb-positive cIN differentiation and maturation can be influenced by sensory input, neuronal activity, or molecular signals (***Dehorter et al., 2015***; ***Hensch, 2005***; ***Huang et al., 1999***; ***Porter et al., 1999***). Our findings align with this concept: Pvalb and other markers of this cIN subtype were rapidly upregulated in mouse MGE cells after grafting onto human organoids, occurring weeks earlier than expected based on *in vivo* timelines (***Figure 4*** and ***Figure 5***). These results suggest that neuronal differentiation and maturation may be accelerated in certain contexts (***Hergenreder et al., 2024***; ***Wallace and Pollen, 2024***), at least for this particular cIN subtype. Chimeric models, such as the ones we described here, offer powerful tools for studying the non-cell-autonomous regulation of neuronal development and maturation, with implications for understanding neurodevelopmental processes, evolution, and disease (***Mostajo-Radji et al., 2020***). The *in vitro* generation of several neuronal subtypes, besides Pvalb cINs, has proven challenging.

While cortical organoids contain radial glia, which theoretically should generate all PN subtypes, specific neurons like layer 4 granular neurons and layer 5 Von Economo and Betz cells have yet to be observed in these models (***Pollen et al., 2019***; ***Quadrato et al., 2017***; ***Velasco et al., 2019***). The differentiation of layer 4 granular neurons seems to rely on thalamic input (***Jabaudon, 2017***; ***Monko et al., 2022***; ***Pouchelon et al., 2014***), and similarly, mature excitatory input from PNs is believed to regulate Pvalb expression in cINs (***Miyamae et al., 2017***; ***Southwell et al., 2012***; ***Wong et al., 2018***). Interestingly, in our chimeric cultures, we observed a significant upregulation of Pvalb in grafted mouse cINs despite the immaturity of the host human PNs, with this upregulation occurring well before mature electrophysiological activity was detected in either human (***Samarasinghe et al., 2021***; ***Sharf et al., 2022***; ***Trujillo et al., 2019***) or mouse (***Ciarpella et al., 2021***; ***Eiraku et al., 2008***; ***Elliott et al., 2023***; ***Park et al., 2024***) 3D *in vitro* cortical models. This suggests that mature PN electrophysiological activity likely did not drive Pvalb induction in these experiments. Moreover, it seems improbable that diffusible factors alone triggered Pvalb fate induction, as neither 2D co-culture with human PNs nor conditioned media exposure resulted in significant Pvalb upregulation (*Fig**ure 6***). Interestingly, we noted a reduced proportion of Sst-positive cINs in co-culture with mouse PNs, compared to those cultured with human PNs or exposed to conditioned media, indicating that diffusible factors may influence Sst identity (***Figure 6***). However, for Pvalb upregulation, it is more plausible that direct cell-to-cell contact within the 3D structure of human cortical cultures was key to the rapid induction of mouse Pvalb cells. Supporting this idea, research in 3D cerebellar development models, including organoids (***Atamian et al., 2024***), and grafting IPSC-derived neurons into human fetal cerebellar slices (***Wang et al., 2015***) has demonstrated that a human 3D architecture is crucial for the proper development and maturation of Purkinje cells, which have long presented a challenge for *in vitro* modeling.

## Limitations

The chimeric models used in this study, including primary organotypic cultures and brain organoids, provide valuable insights into cIN differentiation. However, these *in vitro* systems are inherently artificial and do not fully replicate the complexity of the *in vivo* brain. Furthermore, we previously demonstrated that cellular stress within brain organoids can impair neuronal differentiation (***Bhaduri et al., 2020***). Recent machine learning analyses have also shown that such impairment predominantly affects the transition from radial glia to early PNs, while ventral precursors and cINs in organoids exhibit relatively higher fidelity (***Gonzalez-Ferrer et al., 2024***). Notably, label transfer from primary fetal tissue to organoids has accurately assigned cIN fate across multiple organoid lines (***Gonzalez-Ferrer et al., 2024***).

An alternative hypothesis for the enrichment of Pvalb-positive cINs in our chimeric models is that it may result from selective apoptosis of Sst-positive cINs. In a separate study, *Fezf2* knockout mice demonstrated that Sst-positive cINs undergo selective cell death in response to the absence of subcerebral PNs (***Wu et al., 2024***). However, several lines of evidence suggest that selective cell death is unlikely to account for the Pvalb-positive enrichment observed in our models. First, host human organotypic cultures and organoids still contain Sst-positive cINs that coexist with grafted mouse cINs (***Figure 2***B). Second, in our experiments involving the grafting of lineage-traced Sst cINs, approximately 50% of the cells retained the Sst marker by 14 DPG (***Figure 8***C). Finally, in our grafted cIN progenitors, we observed Pvalb upregulation as early as 2 DPG (***Figure 4***A), indicating that the enrichment of Pvalb-positive cells is more likely driven by the selective induction of this identity rather than the loss of Sst-positive cells.

## Conclusions

In summary, we present a novel chimeric cortical organoid model that accelerates mouse cIN differentiation and significantly increases the proportion of Pvalb-expressing cells under *in vitro* conditions. The specific factors driving this rapid differentiation of mouse MGE cells into Pvalb cINs within human cortical tissue remain unknown. Furthermore, the optimal co-culture conditions required for the efficient differentiation of human Pvalb cINs have yet to be identified. Nonetheless, the chimeric culture system described here provides a robust platform for investigating the mechanisms underlying cIN differentiation and maturation.

## Materials and Methods

### Live primary human cortical tissue collection

All primary tissues were collected and processed under UCSF Gamete, Embryo, and Stem Cell Research Committee (GESCR) approval (Protocol # 10–05113). Patient consent was obtained for all samples, and collection adhered strictly to legal and institutional ethical guidelines. All tissue samples were de-identified, and no sex information was recorded.

Second-trimester samples were obtained from surgical procedures. In compliance with legal and ethical regulations, investigators did not interfere with the surgery. Brain tissue was immediately transported in ice-cold artificial cerebrospinal fluid (ACSF) composed of 125 mM sodium chloride (Millipore Sigma # S9888), 2.5 mM potassium chloride (Millipore Sigma # P3911), 1 mM magnesium chloride (Millipore Sigma # M8266), 2 mM calcium chloride (Millipore Sigma # C4901), 1.25 mM sodium phosphate monobasic (Millipore Sigma # 71505), 25 mM sodium bicarbonate (Millipore Sigma # S5761), and 25 mM D-(+)-glucose (Millipore Sigma # G8270). Before use, this solution was bubbled with 95% O_2_ and 5% CO_2_. All tissue was used within 5 hours of collection.

Upon arrival at the lab, meninges were removed, and the tissue was either processed for organotypic culture or dissociated into cells, as detailed below. Occasionally, tissue with distinguishable medial ganglionic eminences (MGEs) was dissected, sectioned, and cryopreserved for future experiments. Freezing media included Dulbecco’s Modified Eagle Medium: Nutrient Mixture F-12 with GlutaMAX (Thermo Fisher Scientific # 10565018), N-2 Supplement (Thermo Fisher Scientific #17502048), B-27 Supplement (Thermo Fisher Scientific # 17504044), 20 ng/mL Human FGF-basic (Thermo Fisher Scientific # PHG0261), 20 ng/mL Recombinant Human EGF (R&D Systems # 236-EG), 20 µg/mL Human Insulin (Santa Cruz Biotechnology # sc-360248), 5 ng/mL human BDNF (Millipore Sigma # SRP3014), 10 µM Rho Kinase Inhibitor (Y-27632) (Tocris # 1254), 1.6 g/L D-(+)-glucose (Millipore Sigma # G8270), and 10% Dimethyl Sulfoxide (Cell Signaling Technology # 12611).

### Postmortem primary human cortical tissue collection

Deidentified human specimens were collected postmortem with prior patient consent, following the ethical guidelines of the UCSF Committee on Human Research (Protocol # 10-02693). The postmortem interval (PMI) was less than 24 hours. Specimens were evaluated by a neuropathologist and categorized as control samples. The tissue was coronally sectioned, and when possible, one hemisphere was cut axially. One-millimeter blocks were fixed in 4% paraformaldehyde (PFA) (Thermo Fisher Scientific # 28908) for two days, then cryoprotected in a 30% sucrose gradient (Millipore Sigma # S8501). The tissue was embedded in Tissue-Tek O.C.T. Compound (Sakura # 4583), cryosectioned at 30 µm using a cryostat (Leica Biosystems # CM3050), and mounted on glass slides.

### Mouse lines and husbandry

All mouse experiments were conducted in compliance with UCSF IACUC guidelines. The mouse lines used were previously described: *Nkx2.1*-Cre (C57BL/6J-Tg(Nkx2-1-cre)2Sand/J; The Jackson Laboratory # 008661; RRID: IMSR_JAX:008661;(***Xu et al., 2008***)), *Sst*-Cre (Ssttm2.1(cre)Zjh/J; The Jackson Laboratory # 13044; RRID: IMSR_JAX:013044; (***Taniguchi et al., 2011***)), *Pvalb*-Cre (B6.129P2-Pvalbtm1(cre)Arbr/J; The Jackson Laboratory # 017320; RRID: IMSR_JAX:017320; (***Hippenmeyer et al., 2005***)), Ai14 (B6;129S6-Gt(ROSA)26Sortm14(CAG-tdTomato)Hze/J; The Jackson Laboratory # 007908; RRID: IMSR_JAX:007908; (***Madisen et al., 2010***)), Ai34 (B6;129S-Gt(ROSA)26Sortm34.1(CAG-Syp/tdTomato)Hz The Jackson Laboratory # 012570; RRID: IMSR_JAX:012570; (***Daigle et al., 2018***)), and Ai96 (B6J.Cg-Gt(ROSA)26Sortm96(CAG-GCaMP6s)Hze/MwarJ; The Jackson Laboratory # 028866; RRID: IMSR_JAX:028866; (***Madisen et al., 2015***)). Mice were of the C57BL/6J background, and developmental staging was calculated with the plug date as E0.5.

### Pluripotent stem cells maintenance

We used three previously described iPSC lines: H2B.-1323.4 (female; RRID: CVCL_0G84) (***Matsumoto et al., 2013***), H28126 (male) (***Romero et al., 2015***), and WTC-11 (male, Asian; RRID: CVCL_Y803) (***Miyaoka et al., 2014***). Additionally, we employed the human embryonic stem cell (hESC) line HES-3 NKX2.1GFP/w (female; RRID: CVCL_A5HB) (***Goulburn et al., 2011***). Pluripotent stem cells were cultured in StemFlex Medium (Thermo Fisher Scientific # A3349401) with Penicillin/Streptomycin (Thermo Fisher Scientific # 15140122). Cell passaging was done using ReLeSR (Stem Cell Technologies # 05872), following the manufacturer’s instructions. Cells were cryopreserved in mFreSR medium (Stem Cell Technologies # 05855).

### Culture of human and mouse organotypic cortical sections

#### Preparation of ACSF and tissue collection

Fresh ACSF was prepared prior to tissue collection. The composition of ACSF included 125 mM sodium chloride (Millipore Sigma # S9888), 2.5 mM potassium chloride (Millipore Sigma # P3911), 1 mM magnesium chloride (Millipore Sigma # M8266), 2 mM calcium chloride (Millipore Sigma # C4901), 1.25 mM sodium phosphate monobasic (Millipore Sigma # 71505), 25 mM sodium bicarbonate (Millipore Sigma # S5761), and 25 mM D-(+)-glucose (Millipore Sigma # G8270). The solution was continuously bubbled with a gas mixture of 95% O_2_ and 5% CO_2_ to maintain optimal oxygenation and pH balance.

#### Tissue embedding and sectioning

After collection, tissue was stored in ACSF before embedding in 3% low-gelling agarose (Millipore Sigma # A9414). The embedded tissue was sectioned at 300 µM using a vibratome (Leica).

#### Slice culture at the air-liquid interface

Slices were cultured at the air-liquid interface on hydrophilic polytetrafluoroethylene cell culture inserts (Millipore Sigma # PICM0RG50) in a medium containing 32% Hanks’ Balanced Salt Solution (Millipore Sigma # H8264), 60% Basal Medium Eagle (Millipore Sigma # B9638), 5% Fetal Bovine Serum (Millipore Sigma # F2442), 1% D-(+)-glucose (Millipore Sigma # G8270), 1X N2 Supplement, and 100 U/mL Penicillin/Streptomycin. The medium was replenished every 2-3 days to maintain slice viability.

### Human cortical organoids generation

#### iPSC aggregation and early differentiation

To generate human cortical organoids, we followed a modified version of our previously established protocol (Pollen et al., 2019). iPSCs were dissociated into single cells and reaggregated at a density of 10,000 cells per aggregate in lipidure-coated 96-well V-bottom plates, using 100 µL of StemFlex Medium supplemented with 10 µM Rho Kinase Inhibitor (Y-27632, Tocris # 1254) on Day −1.

#### Cortical induction

On Day 0, the medium was replaced with cortical differentiation medium containing Glasgow Minimum Essential Medium (Thermo Fisher Scientific # 11710035), 20% Knockout Serum Replacement (Thermo Fisher Scientific # 10828028), 0.1 mM MEM Non-Essential Amino Acids (Thermo Fisher Scientific # 11140050), 1 mM Sodium Pyruvate (Millipore Sigma # S8636), 0.1 mM 2-Mercaptoethanol (Millipore Sigma # M3148), and 100 U/mL Penicillin/Streptomycin. The medium was further supplemented with 20 µM Rho Kinase Inhibitor (Y-27632, Days 0-6), 3 µM WNT Inhibitor (IWR1-*ϵ*, Cayman Chemical # 13659, Days 0-18), and 5 µM TGF-Beta Inhibitor (SB431542, Tocris # 1614, Days 0-18). Media changes occurred on Days 3, 6, and subsequently every 2-3 days until Day 18. Neuronal differentiation and maintenance: On Day 18, organoids were transferred to ultra-low adhesion plates (Millipore Sigma # CLS3471) and placed on an orbital shaker at 90 revolutions per minute in neuronal differentiation medium. This medium contained Dulbecco’s Modified Eagle Medium: Nutrient Mixture F-12 with GlutaMAX (Thermo Fisher Scientific # 10565018), 10% (v/v) Hyclone characterized fetal bovine serum (Cytova # SH30071.03), 1X N-2 Supplement (Thermo Fisher Scientific # 17502048), 1X Chemically Defined Lipid Concentrate (Thermo Fisher Scientific # 11905031), and 100 U/mL Penicillin/Streptomycin. Media was changed every 2-3 days.

From Day 35 onward, neuronal differentiation medium was supplemented with 5 µg/mL Heparin sodium salt (Millipore Sigma # H3149) and 0.5% v/v Matrigel Growth Factor Reduced (GFR) Basement Membrane Matrix (Corning # 354230).

For scRNA-seq experiments, we removed fetal bovine serum from this media.

#### Neuronal maturation

From Day 70, organoids were cultured in neuronal maturation media containing BrainPhys Neuronal Medium (Stem Cell Technologies # 05790), 10% (v/v) Hyclone characterized fetal bovine serum, 1X N-2 Supplement, 1X Chemically Defined Lipid Concentrate (Thermo Fisher Scientific # 11905031), 1X B-27 Supplement (Thermo Fisher Scientific # 17504044), 100 U/mL Penicillin/Streptomycin, and 1% v/v Matrigel GFR.

### Human MGE organoids generation

#### ESC aggregation and early differentiation

MGE organoids were generated following a modified version of a previously published protocol (***Birey et al., 2017***). The ESC line HES-3 NKX2.1GFP/w was dissociated and reaggregated at a density of 10,000 cells per well in lipidure-coated 96-well V-bottom plates. The cells were cultured in Stem Flex medium supplemented with Rho Kinase Inhibitor Y-27632 on Day −1. On Day 0, the medium was replaced with Dulbecco’s Modified Eagle Medium: Nutrient Mixture F-12 with Gluta-MAX, containing 20% (v/v) Knockout Serum Replacement, 1 mM MEM Non-Essential Amino Acids, mM 2-Mercaptoethanol, and 100 U/mL Penicillin/Streptomycin. The medium was further supplemented with 5 µM dorsomorphin (Millipore Sigma # P5499), 10 µM TGF-beta inhibitor SB431542, and 10 µM Rho Kinase Inhibitor Y-27632. Medium changes occurred on Days 2 and 4, with the Rho Kinase Inhibitor omitted from the subsequent changes.

#### Neuronal differentiation and maintenance

On Day 6, the organoids were transferred to neuronal differentiation medium, which comprised Neurobasal A medium (Thermo Fisher Scientific # 10888022), 1X B-27 supplement minus Vitamin A (Thermo Fisher Scientific # 12587010), 1X GlutaMAX supplement (Thermo Fisher Scientific # 35050061), and 100 U/mL Penicillin/Streptomycin. The medium was replaced every 2-3 days. On Day 18, organoids were moved to ultra-low adhesion plates and cultured on an orbital shaker at 90 revolutions per minute. Neuronal differentiation medium was supplemented with small molecules as follows: From Days 6 to 11, 20 ng/mL Human Recombinant EGF (R&D Systems # 236-EG), 20 ng/mL Human FGF-basic (FGF-2/bFGF) Recombinant Protein (Thermo Fisher Scientific # PHG0261), and 3 µM WNT inhibitor IWR1-*ϵ* were included. On Days 12-15, the medium also contained 100 nM SHH pathway agonist Smoothened Agonist (SAG) HCl (Selleckchem # S7779) and 100 nM retinoic acid (Millipore Sigma # R2625). From Days 16 to 24, retinoic acid was removed, and 100 nM allopregnanolone (Cayman Chemicals # 16930) was added. After Day 25, no small molecules were included in the medium.

### Human thalamic organoids generation

#### iPSC aggregation and early differentiation

Thalamic organoids were generated following a previously described protocol (***Xiang et al., 2020***). Briefly, iPSCs were clump dissociated into ultra-low attachment 6-well plates and cultured for 8 days in Dulbecco’s Modified Eagle Medium: Nutrient Mixture F-12 with GlutaMAX, supplemented with 15% (v/v) Knockout Serum Replacement, 1 mM MEM Non-Essential Amino Acids, 0.1 mM 2-Mercaptoethanol, 100 U/mL Penicillin/Streptomycin, 10 µM SB431542, 100 nM LDN193189 dihydrochloride (Tocris # 6053), and 4 µg/mL human insulin (Santa Cruz Biotechnology # sc-360248). The medium was replaced every other day.

#### Neuronal induction

On day 9, the plates were placed on a shaker operating at 90 rpm, and the organoids were cultured for an additional 8 days in patterning medium, which consisted of Dulbecco’s Modified Eagle Medium: Nutrient Mixture F-12 with GlutaMAX, supplemented with 15% Dextrose (Millipore Sigma # PHR1000), 0.1 mM 2-Mercaptoethanol, 1X N2 Supplement, 2X B-27 Supplement minus Vitamin A, 30 ng/mL recombinant human BMP7 (R&D Systems # 354-BP), 1 µM PD325901 (Millipore Sigma # PZ0162), and 100 U/mL Penicillin/Streptomycin. The medium was replaced every other day.

#### Neuronal differentiation and maintenance

From Day 17 onward, the organoids were cultured in differentiation medium composed of a 1:1 mix of Neurobasal Medium (Thermo Fisher Scientific # 21103049) and Dulbecco’s Modified Eagle Medium: Nutrient Mixture F-12 with GlutaMAX, supplemented with 2X B-27 Supplement, 1X N2 Supplement, 0.1 mM MEM Non-Essential Amino Acids, 100 U/mL Penicillin/Streptomycin, and 50 µM 2-Mercaptoethanol. The medium was replaced every other day.

### Mouse cortical organoids generation

Mouse E14.5 cortices were carefully dissected and chopped into small pieces. The tissue was then dissociated using the Worthington Papain Dissociation System (Worthington # LK003150) according to the manufacturer’s instructions. An enzyme solution was prepared by resuspending 20 units of papain per mL, along with 1 mM L-cysteine and 0.5 mM EDTA in Earle’s Balanced Salt Solution (EBSS). This enzyme solution was activated by incubating at 37°C for 30 minutes. Following activation, 200 units of DNase I per mL were added. The chopped tissue was then transferred into the papain and DNase I solution and incubated at 37°C while shaking at 90 rpm for an additional 30 minutes. The tissue was mechanically dissociated using flamed glass Pasteur pipets (Fisher Scientific # 13-678-6B) and washed in 1X PBS containing 0.1% Bovine Serum Albumin (Millipore Sigma A3311), followed by centrifugation at 300 rcf for 3 minutes.

The resulting cell suspension was resuspended and aggregated at a density of 10,000 cells per well in lipidure-coated 96-well V-bottom plates. The cortical organoid differentiation medium consisted of Dulbecco’s Modified Eagle Medium: 10% (v/v) Hyclone characterized fetal bovine serum, Nutrient Mixture F-12 with GlutaMAX supplement, 1X N-2 Supplement, 1X Chemically Defined Lipid Concentrate, 5 µg/mL Heparin sodium salt, and 1% v/v Matrigel GFR, along with 100 U/mL Penicillin/Streptomycin. The medium was changed every 2-3 days.

After 10 days in culture, the organoids were transferred to neuronal maturation media, which contained BrainPhys Neuronal Medium, 10% (v/v) Hyclone characterized fetal bovine serum, 1X N-2 Supplement, 1X Chemically Defined Lipid Concentrate, 1X B-27 Supplement, 100 U/mL Penicillin/Streptomycin, and 1% v/v Matrigel GFR. At this stage, the organoids were cultured in ultra-low adhesion plates (Millipore Sigma # CLS3471) on an orbital shaker at 90 rpm.

### Dissociation and 2D Culture of Primary Human and Mouse Cortical Neurons

Prior to tissue dissociation, glass-bottom cell culture plates (NEST # 801006) were coated overnight at 37°C with a 0.1% Poly-L-ornithine solution (Millipore Sigma # P4957). The plates were then washed three times with sterile water. Following this, they were coated overnight at 37°C with a mixture of 5 µg/mL Laminin (Millipore Sigma # L2020) and 1 µg/mL Fibronectin (Millipore Sigma # DLW354008) resuspended in PBS.

Mouse E14.5 cortices and human GW22 cortices were dissected and chopped into small pieces. The cortices were then dissociated using the Worthington Papain Dissociation System (for details, see the section on Mouse cortical organoids generation). The resuspended cells were plated at a concentration of 100,000 cells per well. Cells were cultured in cortical organoid differentiation medium, which consisted of Dulbecco’s Modified Eagle Medium: Nutrient Mixture F-12 with GlutaMAX, 10% (v/v) Hyclone characterized fetal bovine serum, 1X N-2 Supplement, 1X Chemically Defined Lipid Concentrate, 5 µg/mL Heparin sodium salt from porcine intestinal mucosa, and 1% v/v Matrigel GFR, along with 100 U/mL Penicillin/Streptomycin. The medium was changed every 2-3 days.

### Dissociation and Infection of Human MGEs

Cryopreserved GW18-19 human MGE tissue stocks were thawed in warm Leibovitz’s L-15 Medium (Thermo Fisher Scientific # 11415064). Following thawing, the cells were dissociated using the Worthington Papain Dissociation System (for details, see the section on Mouse cortical organoids generation). The cells were then concentrated by centrifugation at 300 g for 2 minutes and resuspended in warm cortical differentiation media. The cells were infected for 1 hour with an eGFP adenovirus (Vector Biolabs # 1060), which ubiquitously expresses eGFP under the CMV promoter, at a titer of 1 x 10^7 PFU/ml. After infection, the cells were washed once in warm cortical differentiation media and resuspended in the same media at a concentration of 1,000 cells/µl for immediate grafting onto organoids (for details, see the section on Grafting of MGE-cIN progenitors and Sst-positive cINs). eGFP expression was detectable as early as 12 hours post-infection, with strong expression observed 48 hours post-infection.

### Grafting of MGE-cIN Progenitors and Sst-Positive cINs

E13.5 MGE tissue was microdissected and transferred to ice-cold Leibovitz’s L-15 Medium (Thermo Fisher Scientific # 11415064) supplemented with 180 µg/mL DNase I (Millipore Sigma # 69182). The tissue was mechanically dissociated on ice by pipetting using a P1000 micropipette. Dissociated cells were concentrated by centrifugation for 4 minutes at 300 rcf.

For organoid grafting, the organoids were transferred to individual wells of lipidure-coated 96-well V-bottom plates. A total of 200 µL of fresh cortical organoid neuronal differentiation medium was added, consisting of Dulbecco’s Modified Eagle Medium: Nutrient Mixture F-12 with GlutaMAX, 10% (v/v) Hyclone characterized fetal bovine serum, 1X N-2 Supplement, 1X Chemically Defined Lipid Concentrate (Thermo Fisher Scientific # 11905031), 5 µg/mL Heparin sodium salt from porcine intestinal mucosa, and 1% v/v Matrigel GFR, along with 100 U/mL Penicillin/Streptomycin. Human organoids were 6-8 weeks old at the time of grafting, while mouse organoids had been cultured for 2 weeks prior. We then added 50,000 MGE cells to each well and incubated the organoids for 24 hours at 37°C. Following this incubation, the organoids were carefully transferred to ultra-low-attachment tissue culture plates and incubated on an orbital shaker at 90 rpm at 37°C. The medium was changed every 2-3 days.

For grafting onto human and mouse organotypic cultures, cells were resuspended at a concentration of 1,000 cells/µL. The cell concentrate was then pipetted directly onto the organotypic cultures to facilitate integration.

For co-culture with 2D cortical neurons, 1,000 cells were added to each well of the cortical cultures.

For grafting postmitotic neurons, all ganglionic eminences of *Sst*-Cre::Ai14 mice at E14.5 were microdissected and dissociated. During microdissection, efforts were made to remove the ventricular zone as thoroughly as possible.

### mTOR Inhibition

All experiments utilized 250 nM rapamycin (Millipore Sigma # R8781). For the experimental organoids, rapamycin was incorporated into the media from the time of grafting throughout the experiment. Control experiments received media without rapamycin. The medium was changed every 2-3 days.

### Immunohistochemistry and confocal imaging

Organoids were collected and fixed in 4% paraformaldehyde (PFA) (Thermo Fisher Scientific # 28908) and cryopreserved in 30% sucrose (Millipore Sigma # S8501). They were embedded in a solution containing 50% Tissue-Tek O.C.T. Compound (Sakura # 4583) and 50% 30% sucrose dissolved in 1X phosphate-buffered saline (PBS) at pH 7.4 (Thermo Fisher Scientific # 70011044). Sections were cut to 12 µm using a cryostat (Leica Biosystems # CM3050) and mounted directly onto glass slides. Following three washes of 10 minutes in 1X PBS, the sections were incubated in a blocking solution consisting of 5% v/v donkey serum (Millipore Sigma # D9663), 2% w/v gelatin (Millipore Sigma # G9391), and 0.1% Triton X-100 (Millipore Sigma # X100) for 1 hour. Primary antibodies were incubated overnight at 4°C, followed by three washes for 30 minutes and incubation with secondary antibodies for 90 minutes at room temperature. The sections were then washed three times for 30 minutes in PBS, followed by a final wash in sterile water for 10 minutes. Whole human and mouse organotypic sections were fixed with 4% PFA for 2 hours at room temperature and subsequently washed in PBS at 4°C overnight. Blocking was conducted for one day at 4°C. Primary antibody incubation occurred for 3 days at 4°C, followed by three washes in PBS, each lasting 2 hours. Secondary antibody incubation was also for 3 days at 4°C, followed by similar washing procedures. For human postmortem tissue, mounted slides were defrosted at 4°C for 24 hours and then equilibrated to room temperature for 3 hours. Antigen retrieval was performed using a solution containing 10 mM trisodium citrate dihydrate (Millipore Sigma # S1804) and 0.05% Tween 20 (Millipore Sigma # P1379) at pH 6.0, heated to 95°C for 10 minutes. Samples were washed with TNT buffer (100 mM Tris-HCl [Thermo Fisher Scientific # 15567027], 150 mM sodium chloride [Millipore Sigma # S9888], and 0.1% Tween 20) for 10 minutes, repeated three times, then incubated with 1% hydrogen peroxide (Millipore Sigma # H1009) in PBS for 2 hours. Slides were blocked for 2 hours with TNB solution (100 mM Tris-HCl, 150 mM sodium chloride, and 0.36% w/v bovine serum albumin [BSA] [Millipore Sigma # A2153]). Primary antibodies were incubated overnight at 4°C. The following day, slides were washed three times in TNT buffer. Biotinylated secondary antibodies diluted in TNB were added for 2.5 hours at room temperature. Sections were then incubated with 1:200 streptavidin-horseradish peroxidase (Millipore Sigma # RABHRP3) in TNB for 30 minutes, followed by a 4-minute incubation with tyramide-conjugated fluorophores. The dilutions and order of the used fluorophores were as follows: Cy5 at 1:50. In cases where tyramide was not used, Alexa-conjugated secondary antibodies were employed along with biotinylated secondaries.

Primary antibodies used:

- Rabbit anti-Bdnf (Abcam # ab108319, RRID: AB_10862052; 1:100)
- Rat anti-Bcl11b (Abcam # ab18465, RRID: AB_2064130; 1:100)
- Rabbit anti-Cox6A2 (Novus Biologicals # NBP1-31112; RRID: AB_2085447; 1:100)
- Mouse anti-ErbB4 (Thermo Fisher # MA5-12888; RRID: AB_10986112; 1:100)
- Rabbit anti-Gbx2 (Proteintech # 21639-1-AP; RRID: AB_2878896; 1:250)
- Goat anti-GFP (Abcam # ab6658; RRID: AB_305631; 1:100)
- Mouse anti-HNA (Millipore Sigma # MAB1281; RRID: AB_94090; 1:100)
- Rabbit anti-Map2 (Proteintech # 17490-1-AP; RRID: AB_2137880; 1:100)
- Rabbit anti-MAPK (ERK1/2) phosphorylated (T202/Y204) (Cell Signaling # 9101; RRID: AB_331646; 1:100)
- Rabbit anti-Mef2c (Abcam # ab227085; RRID: AB_3080861; 1:250)
- Mouse anti-Mki67 (BD Biosciences # 550609; RRID: AB_393778; 1:600)
- Rabbit anti-Nkx2.1 (Abcam # ab76013; RRID: AB_1310784; 1:100)
- Mouse anti-Nr2f2 (R&D Systems # PP-H7147-00; RRID: AB_2155627; 1:100)
- Rabbit anti-Pvalb (Swant # PV27; RRID: AB_2631173; 1:250)
- Rabbit anti-Psd95 (Thermo Fisher # 51-6900; RRID: AB_2533914; 1:100)
- Chicken anti-Rbfox3 (NeuN) (Millipore Sigma # ABN91; RRID: AB_11205760; 1:200)
- Chicken anti-RFP (Thermo Fisher Scientific # 600-901-379; RRID: AB_10704808; 1:100)
- Rabbit anti-S6 phosphorylated (pS6) (S235/236) (Cell Signaling # 2211; RRID: AB_331679; 1:100)
- Mouse anti-Satb2 (Abcam # ab51502; RRID: AB_882455; 1:100)
- Mouse anti-Sox2 (Santa Cruz Biotechnology # sc-365823; RRID: AB_10842165; 1:500)
- Mouse anti-Sst (Santa Cruz Biotechnology # sc55565; RRID: AB_831726; 1:100)
- Mouse anti-Tcf7l2 (Millipore Sigma # 05-511; RRID: AB_309772; 1:250)

#### Secondary antibodies

Secondary antibodies were from the Alexa series (Thermo Fisher Scientific), used at a dilution of 1:250. Additionally, biotin-conjugated WFA (Vector Laboratories # B-1355-2; RRID: AB_2336874; 1:200) was visualized using Alexa 488-conjugated streptavidin (Thermo Fisher # S11223; 1:500). Nuclear counterstaining was performed using 300 nM DAPI (4’,6-Diamidino-2-Phenylindole, dihydrochloride) (Thermo Fisher # D1306).

#### Antigen retrieval

Only the Bdnf antibody required antigen retrieval, achieved by incubating the slides at 95°C for 20 minutes in the antigen retrieval solution (10 mM trisodium citrate dihydrate and 0.05% Tween 20 at pH 6.0) prior to blocking.

#### Confocal Imaging

Imaging was conducted using an inverted confocal microscope (Leica CTR 6500) with LAS AF software (Leica). Images were processed using ImageJ software (NIH), while overlays and quantifications were performed using Adobe Photoshop version 2020 (Adobe).

### Whole Organoid Immunostaining, Clearing, and Lightsheet Imaging

Organoids were fixed at room temperature for 45 minutes in 4% paraformaldehyde. After fixation, they were washed three times in PBS and stored at 4°C. For whole-organoid immunostaining and tissue clearing, the organoids were blocked for 24 hours at room temperature in PBS supplemented with 0.2% gelatin and 0.5% Triton X-100 (Millipore Sigma # X100) (PBSGT). Samples were then incubated with primary antibodies for 7 days at 37°C with agitation at 70 rpm in PBSGT supplemented with 1 mg/ml saponin (Quillaja sp; Millipore Sigma # S4521) (PBSGTS).

Following primary antibody incubation, samples were washed six times in PBSGT over the course of one day at room temperature. For nuclear staining, we utilized Syto16 green fluorescent dye (Thermo Fisher Scientific # S7578). Secondary and nuclear staining was performed for 1 day at 37°C with agitation at 70 rpm in PBSGTS. Samples were then washed six times in PBSGT over the course of one day at room temperature.

Whole organoid clearing was performed using ScaleCUBIC-1 solution as described by Susaki et al. (2015). Briefly, the solution contained 25% wt urea (Millipore Sigma # U5378), 25% wt N,N,N’,N’-Tetrakis(2-hydroxypropyl)ethylenediamine (Tokyo Chemical Industry # T0781), and 15% Triton X-100 dissolved in distilled water. Organoids were incubated in ScaleCUBIC-1 solution overnight at room temperature with agitation at 90 rpm. Whole organoid imaging was performed using a custom-made lattice light sheet microscope (UCSF Biological Imaging Development Center), and images were deconvoluted using the Richardson-Lucy algorithm. Image processing was carried out using Imaris 9.2 software (Bitplane).

### EdU (5-Ethynyl-2’-deoxyuridine) labeling

EdU labeling was performed using the Click-iT Plus EdU Cell Proliferation Kit for Imaging® (Thermo Fisher Scientific C10640), adapting the manufacturer’s instructions for tissue slides. Briefly, grafted organoids were treated with 10 µM EdU from the moment of grafting for 48 hours. Organoids were then transferred to media without EdU, and the media was replaced every other day throughout the experiments. After fixation and sectioning (as described in the Immunohistochemistry and confocal imaging section), organoid sections were washed twice for 3 minutes each with 3% bovine serum albumin (BSA) (Millipore Sigma # A2153) dissolved in PBS. The tissue was then permeabilized for 20 minutes using 0.5% Triton X-100 (Millipore Sigma # X100) diluted in PBS, followed by two 3-minute washes in 3% BSA diluted in PBS. Following permeabilization, the tissue was incubated for 30 minutes in a solution containing 88% Click-iT® Reaction Buffer, 2% copper protectant, 0.0024% Alexa Fluor® picolyl azide, and 10% reaction buffer additive. The tissue was then washed twice in 3% BSA. Nuclear counterstaining was performed using 300 nM DAPI.

### Statistical analysis of immunohistochemistry quantifications

Quantifications were performed across 2-5 batches of organoids, with each batch including 3-10 organoids. We analyzed at least 9 sections per organoid, quantifying all cells of interest in each section. For analysis, we summed all cells in each organoid.

All statistical analyses were conducted using Prism 9.3.1 (GraphPad Software). Conditions were compared using an unpaired parametric Student’s t-test without Welch’s correction.

### Single cell transcriptomics analysis of existing datasets

Public datasets from mouse models were analyzed, including E13.5 and E14.5 ganglionic eminences and P10 subcortex (***Mayer et al., 2018***), as well as three samples from the 10X Genomics E18 mouse cortex dataset comprising 1.3 million cells, and E14 and neonatal cortex and subcortex data (***Loo et al., 2019***). These datasets were downloaded as raw FASTQ or BAM files, which were subsequently converted to FASTQ format. Gene quantification was performed using Kallisto (version 0.46.2; https://pachterlab.github.io/kallisto/) (***Bray et al., 2016***) with *Mus musculus* ENSEMBL release 100 transcript annotations.

The Kallisto-Bus output matrix files, encompassing both introns and exons, were processed using CellBender (version 0.2.0; https://github.com/broadinstitute/CellBender) to filter out likely ambient RNA. Only droplets with a probability greater than 0.99 of being cells were retained for analysis. Droplets that detected fewer than 600 genes, or had more than 40% ribosomal or 15% mitochondrial reads, were excluded from the dataset. Doublets were identified and removed using Scrublet (version 0.2.2; https://github.com/swolock/scrublet) with a threshold parameter set to 0.5.

Using Scanpy (version 1.8; https://github.com/theislab/scanpy), counts in droplets were read-depth normalized, log-transformed, and expression values for each gene were scaled across all cells. Principal component analysis (PCA) was performed on the 6000 most variable genes, followed by balanced K-nearest neighbors mapping, UMAP projection, and Leiden clustering (***Polanski et al., 2020***). Clusters exhibiting high expression levels of Gad1, Gad2, and Dlx genes, along with Lhx6 and Nkx2.1, were included in the analysis. The term “VMF” refers to ventromedial forebrain regions, including the septum, preoptic area, and preoptic hypothalamus.

### Fluorescent-Activated Cell Sorting (FACS) Experiments

Grafted organoids were dissociated into single-cell suspensions using the Worthington Papain Dissociation System (Worthington # LK003150) following the manufacturer’s instructions. To minimize RNA degradation, 3,500 units/ml of RNase Inhibitor was added. We processed 10 grafted organoids individually.

After dissociation, the cells were washed twice with PBS and resuspended in PBS supplemented with 1% ultrapure bovine serum albumin (Thermo Fisher Scientific # AM2618) and 4,900 units/ml of RNase Inhibitor. The samples were kept on ice until sorting. FACS was performed using a BD FACSAria Fusion Flow Cytometer with a 130 µm nozzle. Prior to sorting, the samples were incubated with Hoechst 33342 (Thermo Fisher Scientific # 62249). Gating was as follows: Forward and side scatter were used to identify individual cells, while Hoechst 33342 stained nuclei to confirm cell identity. Subsequently, td-Tomato-positive cells were selected for sorting. Each dissociated organoid was sorted separately, and the sorted cells were pooled for single-cell RNA sequencing.

### Single-Cell RNA Library Preparation and Sequencing

A total of 75,000 FACS-purified td-Tomato-positive events (putative cells) were used for this experiment. Immediately after sorting, libraries were prepared using the 10X Chromium Single Cell 3’ reagent kit V3 (10X Genomics # PN-1000092). Following library preparation, the libraries were sequenced on Illumina HiSeq and NovaSeq platforms.

### Single-cell RNA sequencing data processing

Transcriptomes were aligned to both the human (GRCh38) and mouse (mm10) reference genomes, integrating the td-Tomato sequence into the mouse genome. Alignments were performed using Cell Ranger version 5 (10X Genomics). Cells aligning to the human genome and td-Tomato-negative mouse cells were excluded, resulting in a final dataset of 225 td-Tomato-positive mouse cells. Subsequent analysis was carried out in Scanpy 1.10.2 (***Wolf et al., 2018***). The raw counts expression matrix was loaded as an AnnData object, normalized by read depth, log-transformed, and scaled per gene across the dataset. Cells with fewer than 200 detected genes and genes expressed in fewer than three cells were filtered out. The raw and processed data have been deposited in GEO, under accession number GSE278531.

### Deep learning-based cell classification

Td-Tomato-positive cells were classified using SIMS (RRID: SCR_025787), a transformer-based label transfer deep learning architecture previously validated on neuronal single-cell data (***Gonzalez-Ferrer et al., 2024***). The SIMS model was trained on a subset of single-cell RNA expression data with pre-annotated labels from a published dataset compilation (***Schmitz et al., 2022***), specifically focusing on MGE and cortical interneurons. This subset included 51,402 cells and 15 cell labels. The trained model is accessible via the SIMS web app: https://sc-sims-app.streamlit.app/.

### Soma Size Measurements

To minimize potential bias in image acquisition, we reused images obtained during marker quantifications, selecting random sections for analysis. Files were processed using ImageJ 2.3.0. We utilized the Z Project function to create a single-plane image from the z-stacks, selecting Maximal Intensity as the projection type. The freehand selection tool was then employed to delineate the area of the maximal soma size. Finally, the Measure tool was used to calculate the area of the largest soma.

### Live Imaging of Organoids

Live imaging was conducted as previously described (***Huang et al., 2020***). Initially, grafted organoids were transferred to hydrophilic polytetrafluoroethylene cell culture inserts (Millipore Sigma # PICM0RG50) and placed on a six-well glass-bottom tissue culture plate to facilitate culture in an air-liquid interface. The medium for culture was cortical organoid neuronal differentiation medium, excluding Matrigel. Organoids were incubated for 6 hours at 37°C prior to imaging to allow for tissue flattening.

Imaging was performed on an inverted Leica TCS SP5 confocal microscope equipped with an on-stage incubator that maintained a chamber atmosphere of 5% CO_2_, 8% O_2_, and balanced N_2_. The chamber temperature was set to 37°C. Slices were imaged continuously for 24 hours using a 10X air objective (with 2X zoom) at 20-minute intervals.

### Calcium Imaging

Calcium imaging was carried out using the genetically encoded calcium indicator GCaMP6s, specifically recombined in the cINs. Imaging was performed on an inverted confocal microscope (Leica CTR 6500) utilizing LAS AF software (Leica). Images were captured every 1.2 seconds and subsequently processed using ImageJ software.

## Acknowledgments

We are grateful to the families who generously donated tissue samples. We thank Tomasz Nowakowski and David Haussler for their feedback and discussions throughout the development of this manuscript, Daniel Lim for providing laboratory space to carry out part of this research, David Shin for supplying the thalamic organoids, Sainath Mamde and Sara Medor for their expertise in bioinformatics, Steven Cincotta for contributing mouse tissue, and Arpana Arjun and Kevin Barber for their assistance with calcium imaging. We also acknowledge Kyle Marchuk and the UCSF Biological Imaging Development Center for their support in light-sheet imaging. We thank Alain Chedotal and his team for generously sharing protocols for thick tissue immunostaining and clearing, which we accessed through the Tissue Clearing and 3D Imaging Course at the Institut de la Vision (Paris, France). Additionally, we are grateful to Andrew Elefanty for sharing the hES-3 NKX2.1-GFP reporter line.

This research was supported by Schmidt Futures (SF857) awarded to M.T. and A.A.P., the Chan Zuckerberg Biohub to A.A.P., the UCSF Program for Breakthrough Biomedical Research (PBBR) to A.A.P., the National Human Genome Research Institute (1RM1HG011543) to M.T., the National Science Foundation (NSF grants NSF2134955 and NSF2034037) to M.T., and the National Institute of Mental Health (1U24MH132628) to M.A.M.-R. M.A.M.-R. received partial support from the TL1 TR001871 fellowship from the NIH National Center for Advancing Translational Sciences. H.E.S. is supported by the National Science Foundation Graduate Student Research Fellowship.

## Author contributions

M.A.M.-R., A.A.B. and A.A.P. conceived the project. M.A.M.-R., W.R.M.L., A.B., J.G.F., H.E.S., J.L., L.Z., M.T.S., Y.P., T.Z., M.G.A., J.C., F.N.S., D.T., E.C.C. performed the experiments and analyzed the data. M.A.M.-R., M.F.P., M.T., A.R.K., A.A.B. and A.A.P. supervised the work. M.A.M.-R. and A.A.P. wrote this manuscript with contributions from all authors.

