## Supplemental Figures 1-15 for "Fate plasticity of interneuron specification"

Supplemental Figure 7. Validation of Mef2c antibody……………..…………………….8

Supplemental Figure 8. 2D differentiations produce Sst-positive but not Pvalb-positive cINs…......................................................................................................................…...……..9

Supplemental Figure 9. Validation of MGE and thalamic organoid protocols…………....10

Supplemental Figure 10. Human cINs fail to upregulate Pvalb at 7 DPG………………11

Supplemental Figure 11. Immunohistochemistry of 45 days-old human organoids cultured in serum-free conditions…………………………………………………………………12

Supplemental Figure 12. Non-cell-autonomous regulation of Pvalb fate through FBS depletion…………………………………………..………………………………13

Supplemental Figure 13. Fluorescence activated cell sorting (FACS) of grafted organoids grown in FBS-free conditions.………..……………………………………………………15

Supplemental Figure 14. Rapamycin's effect on ERK/MAPK and mTOR pathways.….16

Supplemental Figure 15. Graft of E14.5 mouse Sst cINs onto human organoids……...17

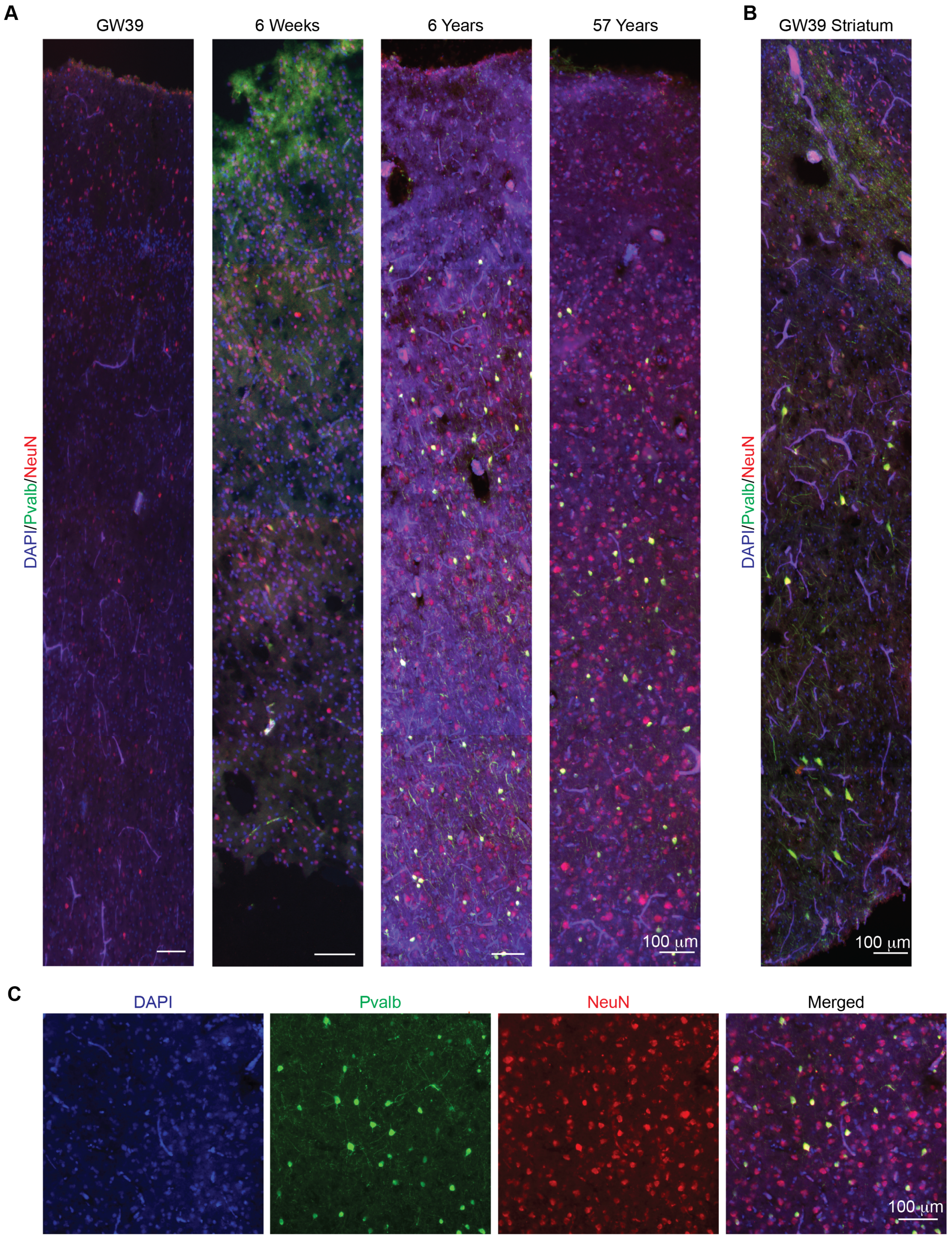

**Supplemental Figure 1. Pvalb immunoreactivity in human postmortem brain samples. A.** Immunohistochemical staining for Pvalb and NeuN in the cerebral cortex of donors across different developmental stages: gestational week 39 (GW39), 6 weeks post-birth, 6 years, and 57 years of age. **B.** Pvalb and NeuN immunostaining in the striatum of the same GW39 donor shown in panel A. **C.** High-magnification image of Pvalb and NeuN immunostaining in the cerebral cortex of a 57-year-old donor.

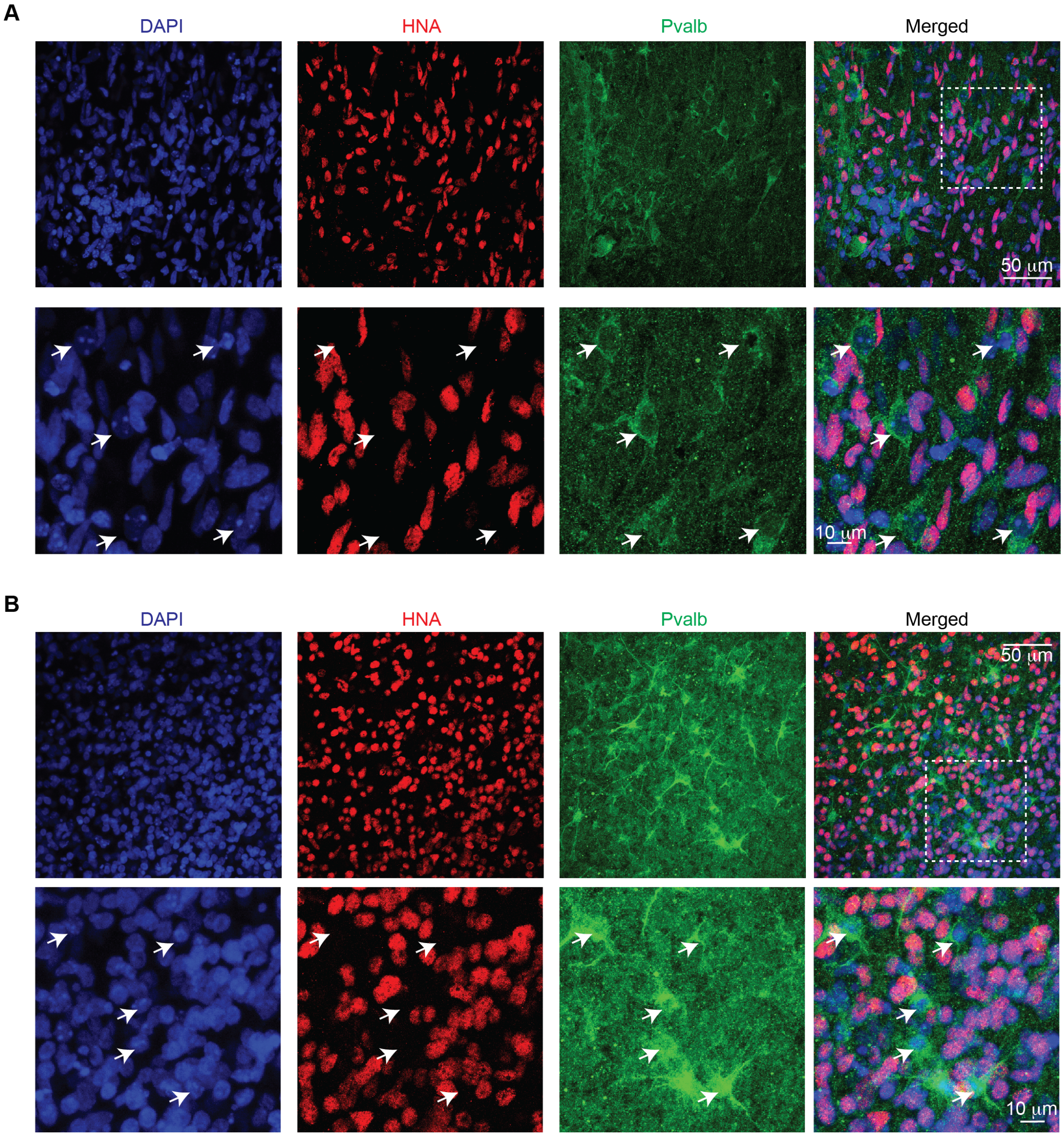

**Supplemental Figure 2: Additional examples of unlabeled grafts.** Immunohistochemical labeling for Human Nuclei Antigen (HNA) and Pvalb in human brain slices grafted with mouse cINs. **A.** Immunostaining in tissue from donor 2. **B.** Immunostaining in tissue from donor 3.

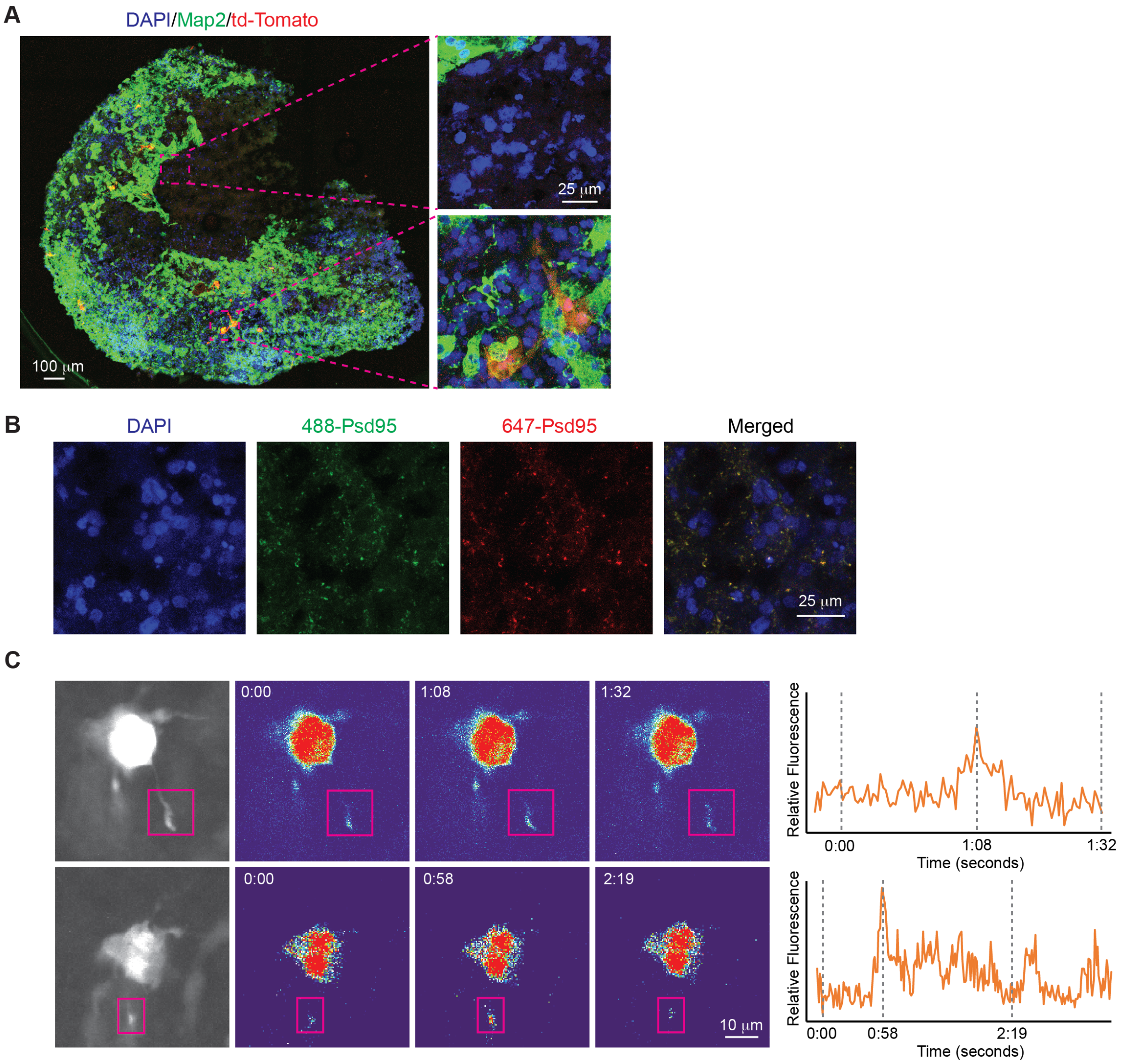

**Supplemental Figure 3. Grafted cINs migrate to neuronal regions of the organoid. A.** The section of an organoid labeled with the pan-neuronal marker Map2 shows that grafted cINs become localized to regions of the organoid containing higher numbers of neurons. **B.** Validation of the Psd95 using two secondary antibodies (Alexa 488 and Alexa 647) show complete colocalization, suggesting true staining. **C.** Examples of calcium transients in grafted cINs 4 MPG. Magenta rectangles = Axodendritic processes with calcium transients.

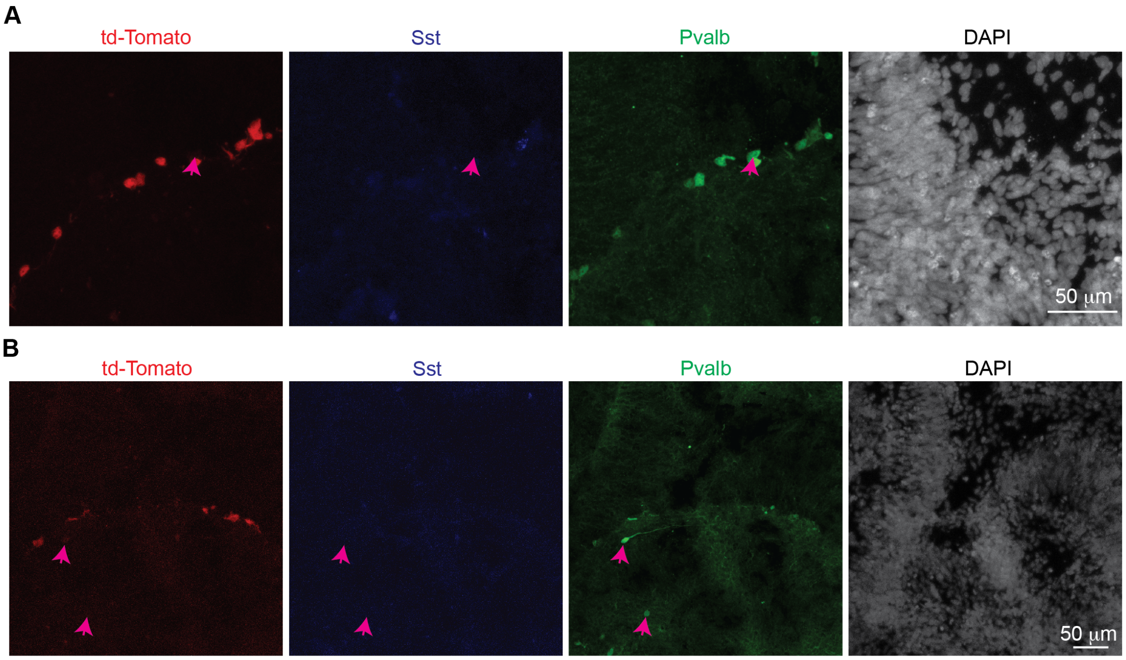

**Supplemental Figure 4. Additional examples of Pvalb immunoreactivity at 2 DPG.**
**A-B.** Immunostaining for Sst and Pvalb in human organoids at 2 DPG. Magenta arrows indicate cells with Pvalb immunoreactivity and low or absent td-Tomato expression.

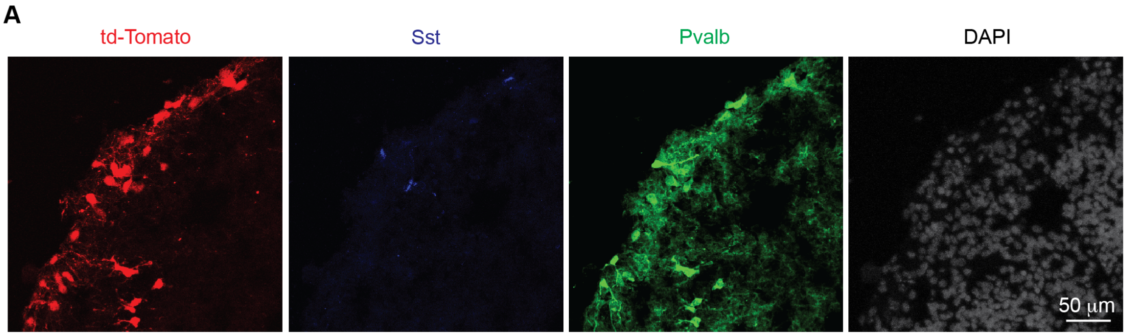

**Supplemental Figure 5. Additional examples of Pvalb immunoreactivity at 7 DPG.**
**A.** Immunostaining for Sst and Pvalb in human organoids at 7 DPG.

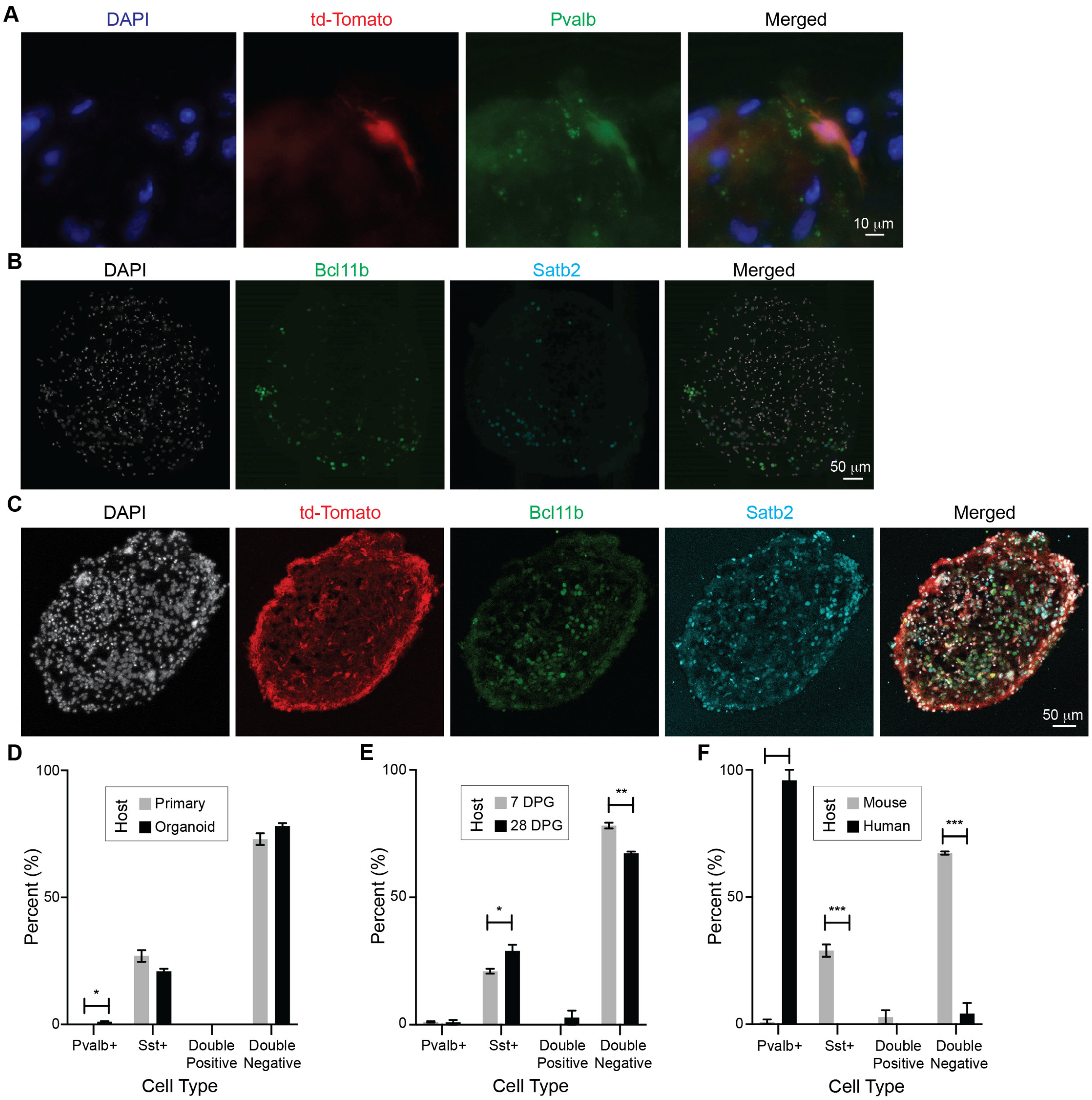

**Supplemental Figure 6. Development of organoid models for cIN grafting. A.** Grafting of MGE progenitors from the Pvalb reporter line results in td-Tomato upregulation at 7 DPG, confirmed by Pvalb immunostaining. **B.** Mouse organoids contain major cortical subtypes, demonstrated by immunostaining for the corticofugal PN marker Bcl11b (Ctip2) and the callosal PN marker Satb2. Grafted cINs are labeled with td-Tomato. **C.** Grafting of mouse cINs onto mouse organoids. **D.** Comparison of cIN subtypes generated at 7 DPG in primary mouse organotypic cultures (as shown in Figures 2C-D) and mouse organoids. **E.** Comparison of cIN subtypes generated at 7 DPG and 28 DPG in mouse organoids. **F.** Comparison of cIN subtypes generated at 28 DPG in mouse versus human organoids. * = $p < 0.05$; ** = $p < 0.01$; *** = $p < 0.001$; **** = $p < 0.0001$. Error bars represent SEM.

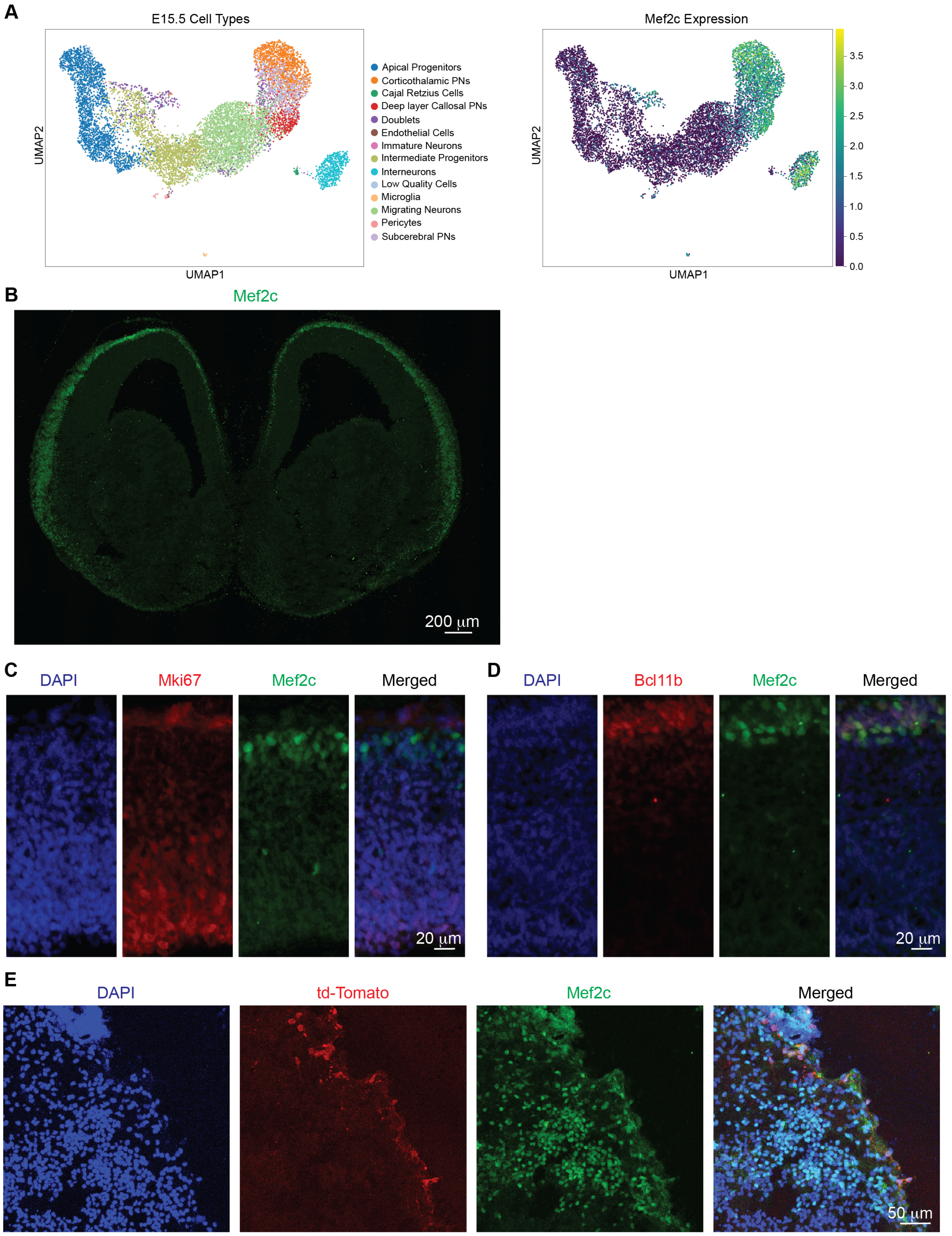

**Supplemental Figure 7. Validation of Mef2c Antibody. A.** Expression of *Mef2c* in the E15.5 mouse cortex. Left: All cell types present in the dataset. Right. Mef2c Expression. Data from Di Bella et al., 2021. **B**. Mef2c immunostaining profile in a coronal section of an E15.5 mouse brain. **C.** Colocalization of Mef2c with the proliferation marker Mki67 shows exclusion of Mef2c from the progenitor-rich ventricular zone. **D.** Colocalization of Mef2c with the corticofugal PN marker Bcl11b shows enrichment of Mef2c in maturing neurons. **E.** Mef2c immunostaining in human organoids grafted with mouse cINs.

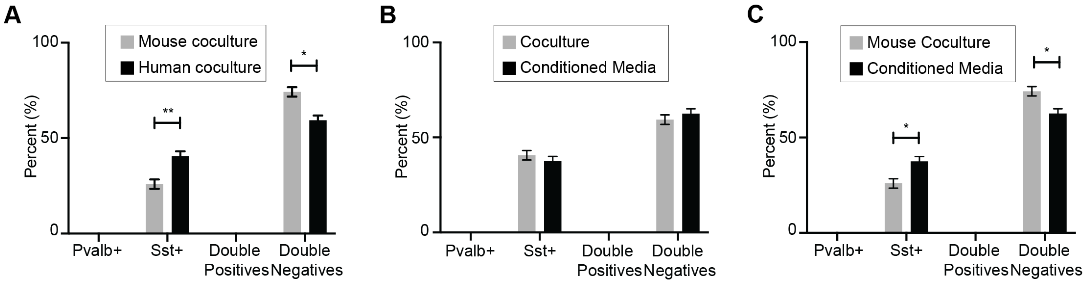

**Supplemental Figure 8. 2D differentiations produce Sst-positive but not Pvalb-positive cINs. A.** Comparison of cIN differentiation in coculture with mouse versus human cortical cells. **B.** Comparison of cIN differentiation in human cortical cell coculture versus media conditioned by human cortical cells. **C.** Comparison of cIN differentiation in mouse cortical cell coculture versus media conditioned by human cortical cells. * = $p < 0.05$; ** = $p < 0.01$. Error bars represent SEM.

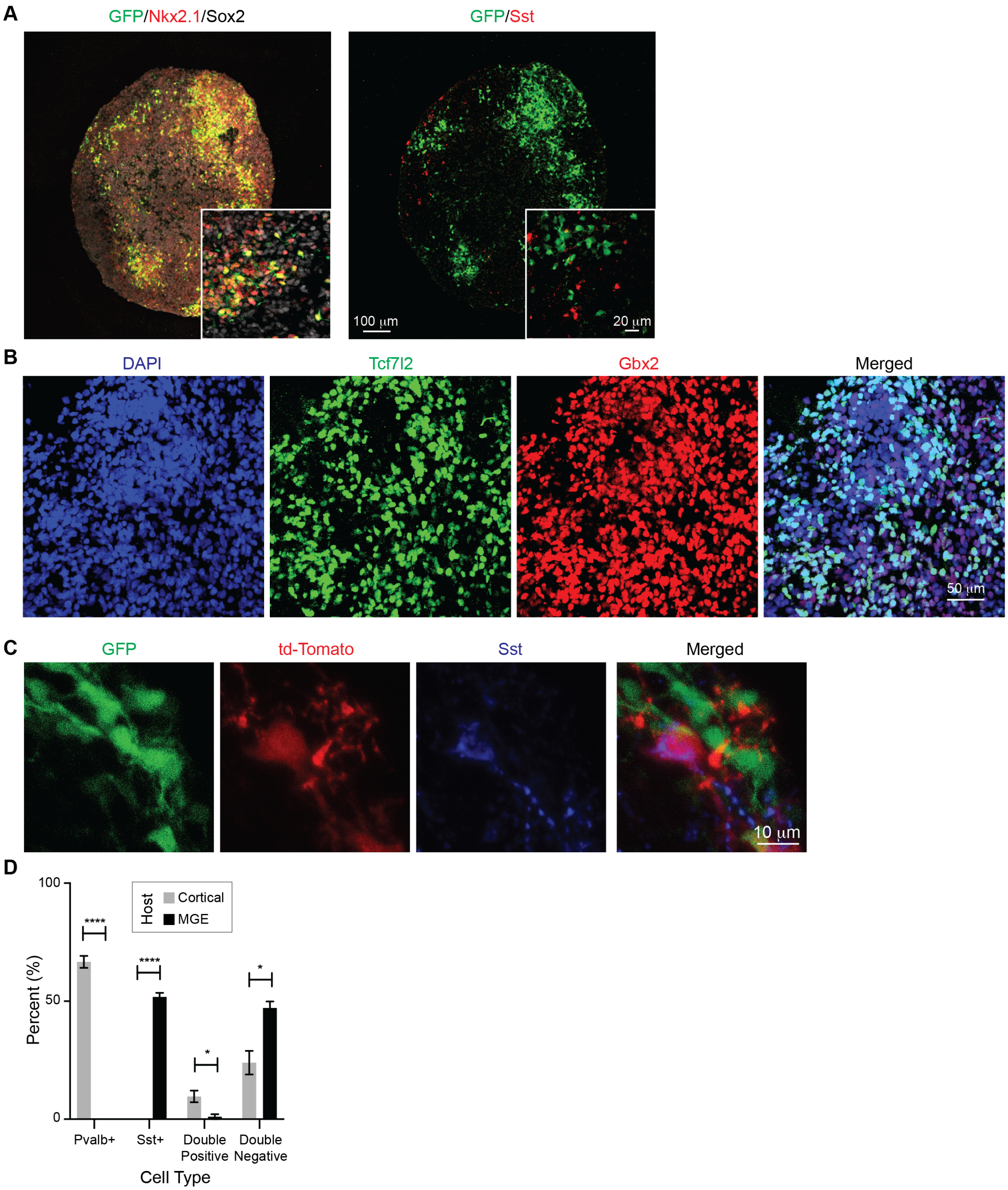

**Supplemental Figure 9. Validation of MGE and thalamic organoid protocols. A.** MGE organoid differentiation was validated using the Nkx2.1-GFP hES cell line, and immunostaining for the progenitor marker Sox2, Nkx2.1, and Sst. Images were taken at 21 DIC. **B.** Thalamic organoid differentiation was validated using immunostaining for thalamic markers Tcf7l2 (also known as Tcf4) and Gbx2. Images were taken at 12 WIC. **C.** Validation of Sst upregulation in the Nkx2.1-GFP hES cell line. **D.** Comparison of cell types generated in human cortical and MGE organoids at 7 DPG. * = $p < 0.05$; **** = $p < 0.0001$. Error bars represent SEM.

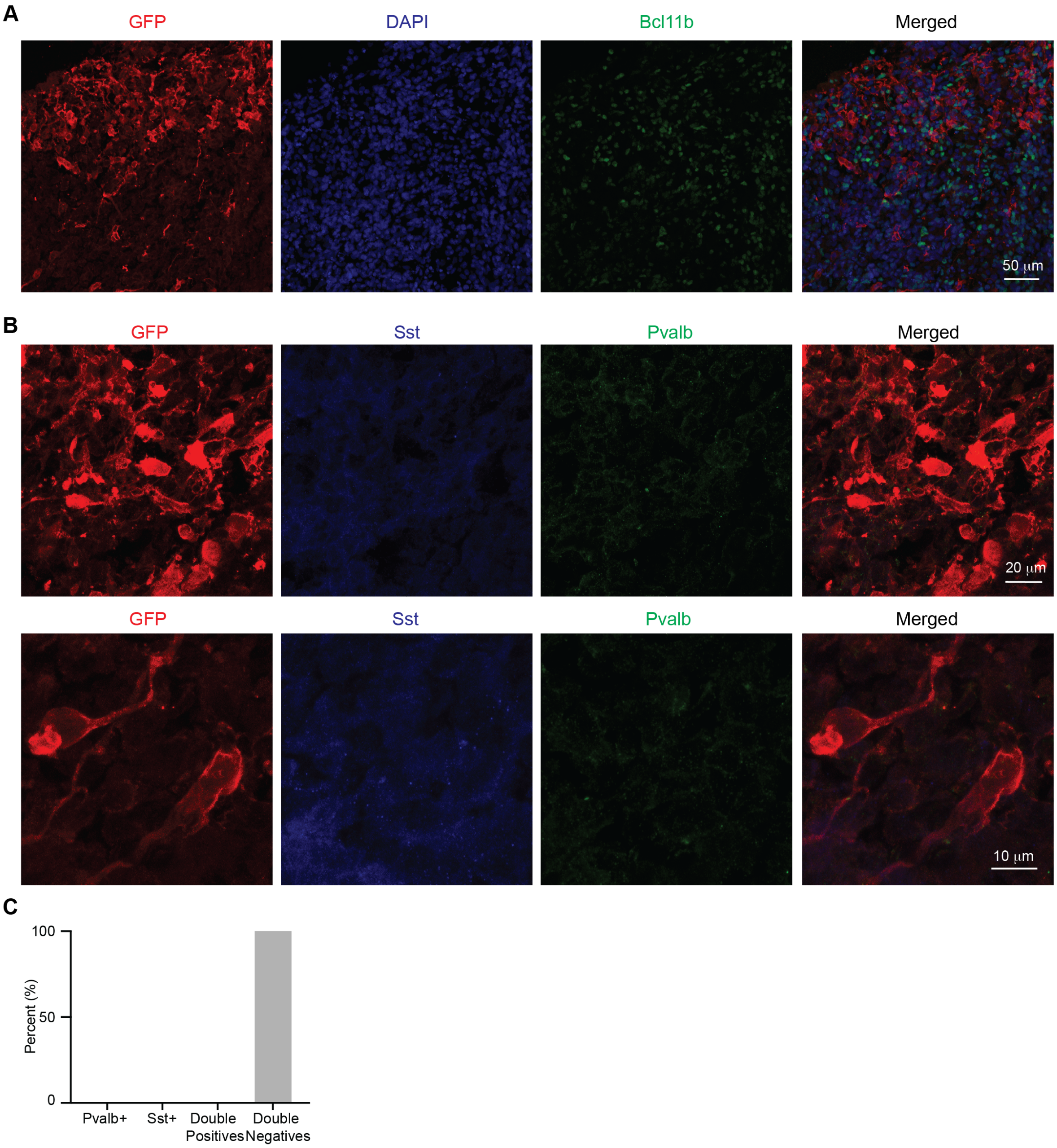

**Supplemental Figure 10. Human cINs fail to upregulate Pvalb at 7 DPG. A.** Representative image of human cINs grafted onto human cortical organoids, with Bcl11b immunostaining indicating the cortical identity of the host organoid. **B.** Representative images of immunostaining for Sst and Pvalb in grafted human cINs. **C.** Quantification of the identity of grafted human cINs at 7 DPG.

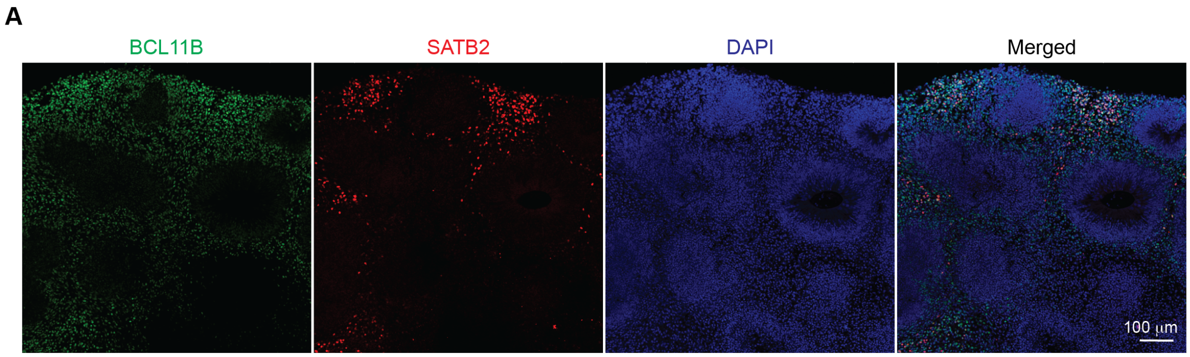

**Supplemental Figure 11. Immunohistochemistry of 6-week-old human organoids cultured in serum-free conditions. A.** Immunostaining showing the corticofugal PN marker BCL11B and the callosal PN marker SATB2.

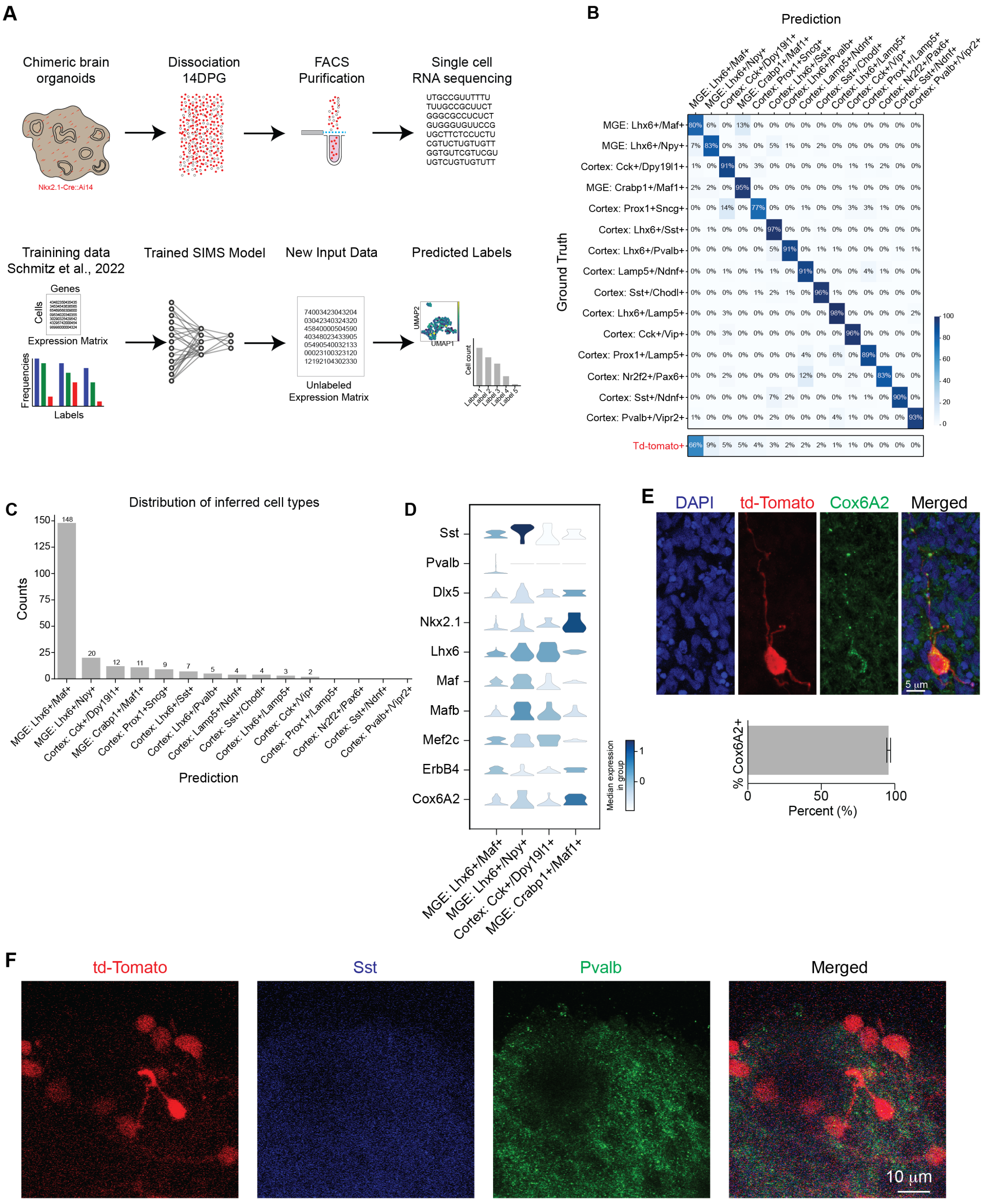

**Supplemental Figure 12. Non-cell-autonomous regulation of Pvalb fate through FBS depletion. A.** Experimental Design: Top: Schematic outlining the experimental approach. Transcriptomic analysis was performed on mouse cINs grafted onto human organoids grown in FBS-free media. The organoids were dissociated at 14 DPG, and fluorescence activated cell sorting (FACS) was used to enrich for td-Tomato-positive cells.

Human cells and td-Tomato-negative cells were bioinformatically excluded, leaving 225 td-Tomato-positive mouse cells for subsequent analysis. Bottom: SIMS, a deep learning model, was trained using 80% of the cells (42,024 out of 51,402 cells) from an atlas of developing cINs, including immature MGE-derived cells and postnatal cINs from the Schmitz et al., 2022 dataset. Label transfer was then applied to the bioinformatically purified td-Tomato-positive cells for cell identity inference. **B.** Confusion Matrix of SIMS Model Predictions: Top: Validation of the SIMS model using the remaining 20% of cells (10,506 cells) from the Schmitz et al., 2022 dataset. The model achieved an accuracy of 89.94% and a Macro F1 score of 0.83. Bottom: SIMS predictions for the td-Tomato-positive cells from the grafted organoids. **C.** Histogram of Inferred Cell Types. The majority (148 out of 225) of the td-Tomato-positive cells were classified into the immature MGE Lhx6+ Maf+ cluster. **D.** Violin plots of marker gene expression profiles in the td-Tomato-positive cells, highlighting cell types represented by more than 10 cells. **E.** Immunostaining and quantification of Cox6A2 in cINs grafted onto human organoids cultured under FBS-free conditions. Error bars represent SEM. **F.** Immunostaining of Pvalb and Sst in cINs grafted onto human organoids under FBS-free conditions.

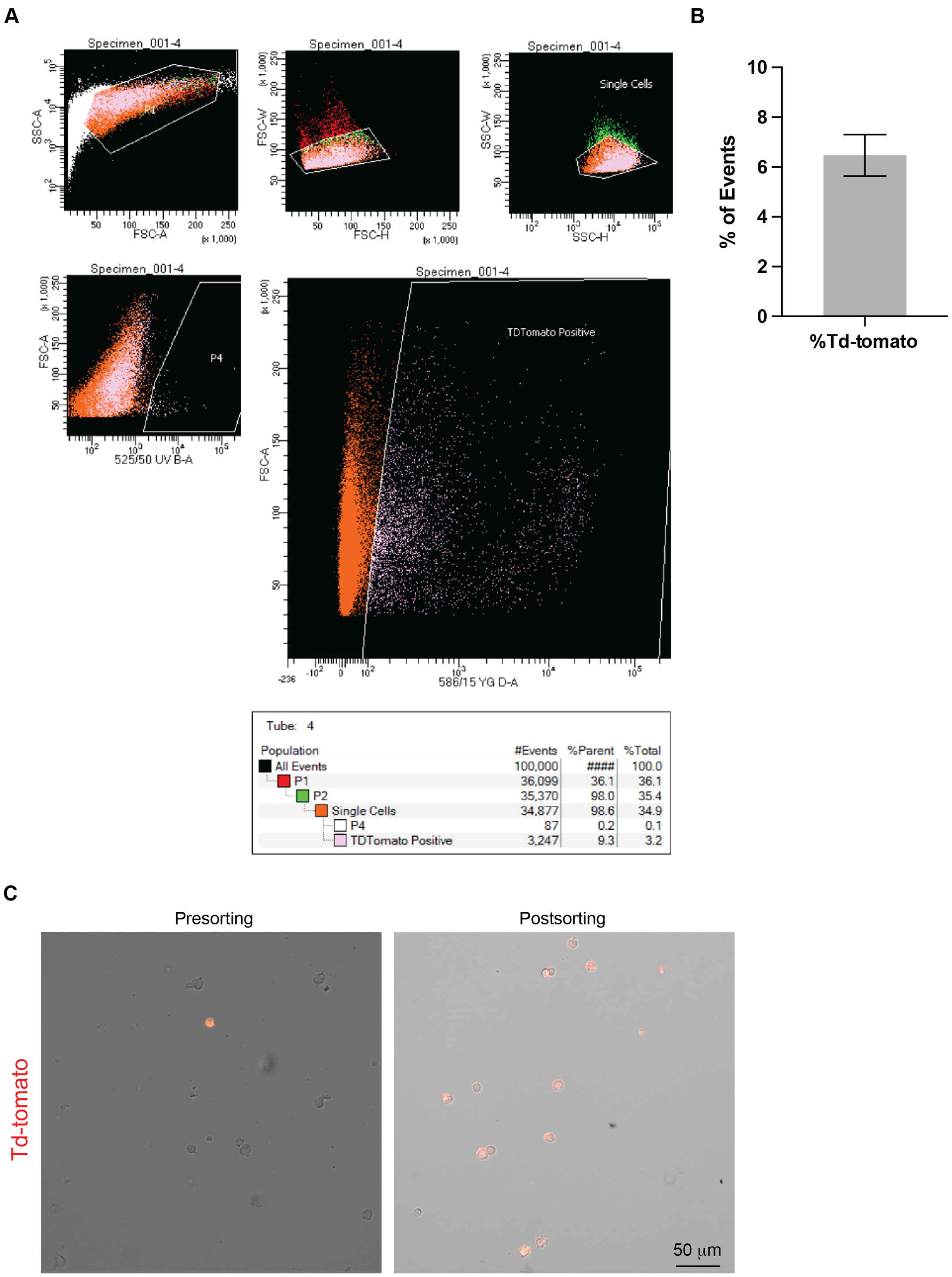

Tube: 4

| Population | #Events | %Parent | %Total |
|---|---|---|---|
| All Events | 100,000 | #### | 100.0 |
| P1 | 36,099 | 36.1 | 36.1 |
| P2 | 35,370 | 98.0 | 35.4 |
| Single Cells | 34,877 | 98.6 | 34.9 |
| P4 | 87 | 0.2 | 0.1 |
| TDTomato Positive | 3,247 | 9.3 | 3.2 |

**Supplemental Figure 13. Fluorescence activated cell sorting (FACS) of grafted organoids grown in FBS-free conditions. A.** Gates setup for the FACS experiment. **B.** Percentage of td-Tomato-positive events observed across all 10 dissociated organoids. Td-tomato-positive events represented 6.47 ± 2.50% of all events. **C.** Representative images showing the dissociated cells before and after FACS sorting.

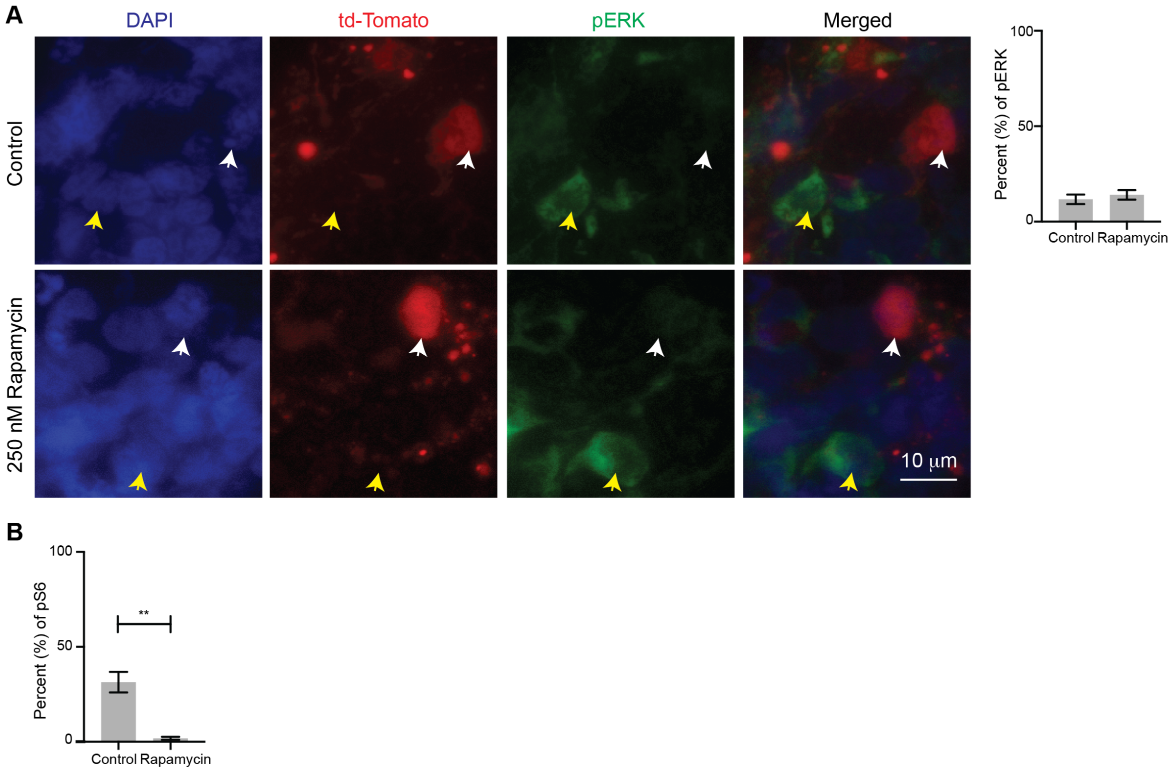

**Supplemental Figure 14. Rapamycin's effect on ERK/MAPK and mTOR pathways. A.** Representative images and quantification of phosphorylated ERK1/2 (pERK) in control versus Rapamycin-treated organoids at 14 DPG. White arrows indicate td-Tomato-positive cells that are negative for pERK, while yellow arrows indicate cells negative for td-Tomato but positive for pERK. **B.** Quantification of phosphorylated ribosomal protein S6 (pS6) in host cells from control and Rapamycin-treated organoids at 14 DPG. ** = $p<0.01$. Error bars represent SEM.

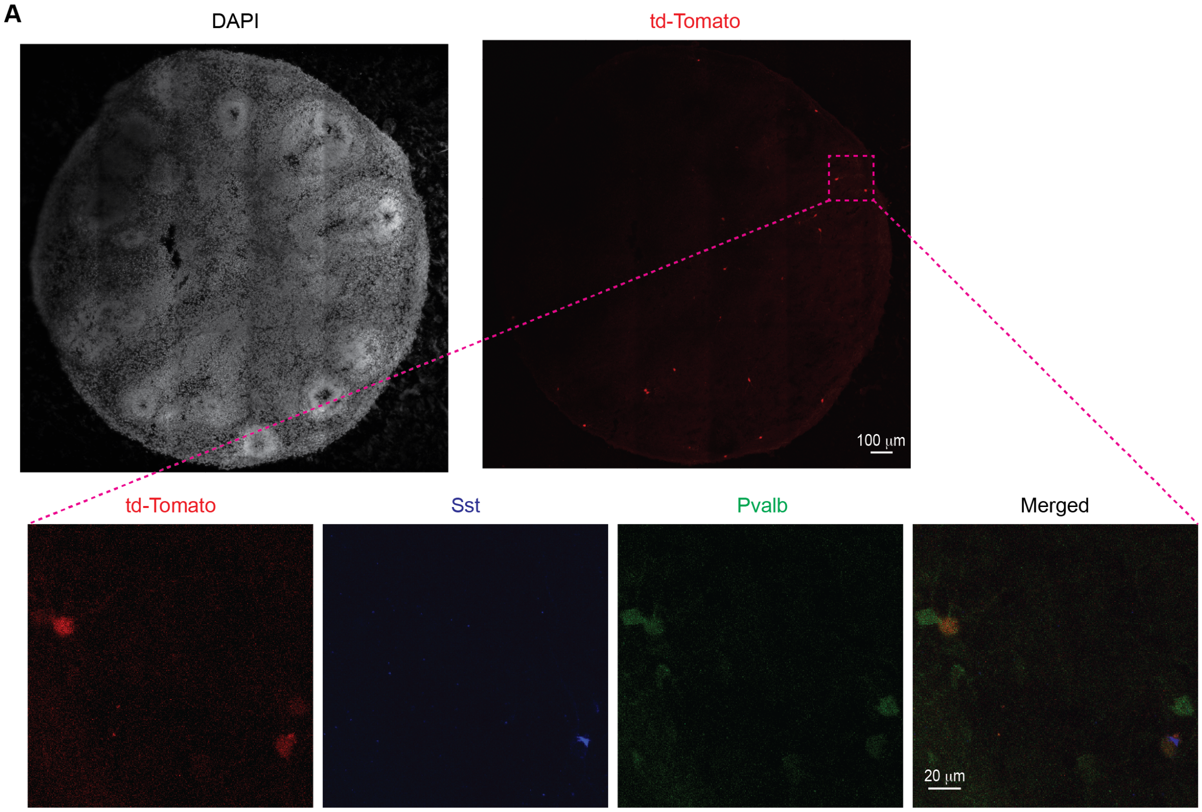

**Supplemental Figure 15. Graft of E14.5 mouse Sst cINs onto human organoids. A.** Top. Whole section image of a grafted organoid showing the td-Tomato recombination in the *Sst*-Cre lineage. Bottom. Example of immunostaining for Sst and Pvalb in *Sst*-Cre::Ai14 grafted organoids.
